# Coessential Genetic Networks Reveal the Organization and Constituents of a Dynamic Cellular Stress Response

**DOI:** 10.1101/847996

**Authors:** David R. Amici, Jasen M. Jackson, Kyle A. Metz, Daniel J. Ansel, Roger S. Smith, Sonia Brockway, Seesha R Takagishi, Shashank Srivastava, Brendan P. O’Hara, Byoung-Kyu Cho, Young Ah Goo, Neil L. Kelleher, Issam Ben-Sahra, Daniel R. Foltz, Marc L. Mendillo

## Abstract

The interrelated programs essential for cellular fitness in the face of stress are critical to understanding tumorigenesis, neurodegeneration, and aging. However, modelling the combinatorial landscape of stresses experienced by diseased cells is challenging, leaving functional relationships within the global stress response network incompletely understood. Here, we leverage genome-scale fitness screening data from 625 cancer cell lines, each representing a unique biological context, to build a network of “coessential” gene relationships centered around master regulators of the response to proteotoxic, oxidative, hypoxic, and genotoxic stress. This approach organizes the stress response into functional modules, identifies genes connecting distinct modules, and reveals mechanisms underlying cellular dependence on individual modules. As an example of the power of this approach, we discover that the previously unannotated HAPSTR (C16orf72) promotes resilience to diverse stressors as a stress-inducible regulator of the E3 ligase HUWE1. Altogether, we present a broadly applicable framework and interactive tool (http://fireworks.mendillolab.org/) to interrogate biological networks using unbiased genetic screens.

## INTRODUCTION

All living organisms retain an evolutionarily conserved capacity to adapt to physiological stress. Highly relevant to human disease, cellular stress response pathways empower aging neurons to clear protein aggregates, sun-exposed melanocytes to repair damaged DNA, and large tumors to proliferate despite having outgrown their blood supply. Certain types of stress, such as thermal stress, are easily studied. Others, however, such as the landscape of misfolded proteins in tumors, are more difficult to model experimentally. Moreover, diseased cells rarely face one stress in isolation. Do cells grown in a hypoxic incubator in pH-buffered, nutrient-rich media adequately model the stresses faced by an ischemic cardiomyocyte after vessel occlusion? Isolated stress perturbations are invaluable to provide mechanistic insight into individual stress response programs, but they fail to capture the complexity that may be found in disease states where these pathways function in a global, integrated network. Unfortunately, it is intractable to comprehensively study all possible combinations of different stresses and stress doses. Alternative strategies are needed to probe stress response pathways in disease-relevant contexts and to identify the mechanisms by which these pathways are integrated.

One approach to investigate context-specific biological relationships is to leverage the heterogeneity of cancer cell lines, each representing a unique genomic and epigenomic landscape obtained through rapid evolution from diverse cells of origin. In studying stress biology, cancer cells may be a particularly suitable model of physiologically relevant states, as tumors routinely co-opt stress response systems to withstand the physiologic challenges imposed by carcinogenesis (Luo et al., 2009). For example, individual tumors may suppress the Hippel-Lindau tumor suppressor (VHL) to induce constitutive hypoxia signaling (Maxwell et al., 1999). Other tumors may alternatively splice pyruvate kinase to promote a metabolic shift toward aerobic glycolysis (Christofk et al., 2008). Still others may activate Heat Shock Factor 1 (HSF1) to coordinate protein homeostasis (proteostasis) during anabolic proliferation (Santagata et al., 2013). Thus, the contextual breadth of cancer cells and the multitude of mechanisms by which they activate stress response pathways may provide unique insights into the regulation and integration of these pathways in diseased cells.

Large-scale fitness screening efforts have now quantified the essentiality of most protein-coding genes across hundreds of different cancer cell lines (Hart et al., 2015; Meyers et al., 2017; Tsherniak et al., 2017). These studies reveal that, even in cancer cells, most critical regulators of the stress response are not universally essential for cell viability. But are all cancer cell lines similarly affected by the loss of individual stress response factors? If not, what mechanisms underlie the requirement for these factors in some cell lines but not others, and which other genes share this context-specific phenotype? Regarding the latter question, recent work has demonstrated that correlated patterns of gene essentiality (”coessentiality”) across many cancer cell lines can indeed identify functional relationships between genes (Boyle et al., 2018; Kim et al., 2019; Pan et al., 2018; Wang et al., 2017). These studies utilize a top-down approach, organizing the genome into coessential clusters which have already proven to yield valuable insights. However, due to technical limitations, such as spurious correlations between genes within genomic regions subject to copy number variation and the relatively modest signal for genes which function outside of obligate protein complexes, much of the genome is left incompletely explored by these methods. Overcoming these technical limitations may thus expand the reach of this approach to biological systems of greater regulatory complexity, such as the cellular stress response network.

In this article, we develop a framework to use genome-scale, CRISPR-Cas9 fitness screening data from 625 cancer cell lines to build a bottom-up network of coessential relationships centered around critical regulators and effectors of the stress response. Our approach organizes functional modules within the stress response, corresponding to both canonical and novel functions of these factors, and identifies genes which represent points of crosstalk between distinct modules. We demonstrate that context, in tumor subtype and cell lineage, is associated with dynamic changes in the organization and utilization of stress response modules in a manner reflecting therapeutically-targetable biology. Finally, we identify novel genes in a global stress network, validating that the previously unannotated HAPSTR (C16orf72) promotes cellular adaptation to diverse stressors as a stress-inducible regulator of HUWE1, an E3 ligase which modulates genotoxic, proteotoxic, and hypoxic stress responses (Kao et al., 2018). More broadly, we demonstrate the power of our approach using the global stress response network as proof of principle and provide an interactive web-based tool (http://fireworks.mendillolab.org/) to facilitate studies of other biological networks.

## RESULTS

### Assembling a coessentiality network around the master regulator of cytosolic proteostasis

Previous reports have demonstrated that genes with shared functions, particularly those which encode components of obligate protein complexes, have convergent effects on fitness when deleted in cancer cell lines (Kim et al., 2019; Pan et al., 2018). Consistent with prior analyses performed in few cell lines, network analysis of the strongest coessential relationships in the genome (r>0.6; 1532 pairings) successfully identifies modules of biological relevance with highly correlated fitness profiles (Figure S1B). However, these clusters are dominated by few pathways, many of which center on major protein complexes – primarily, the ribosome, oxidative phosphorylation, and spliceosome machinery. Indeed, the gene pairs with the most correlated (coessential) knockout fitness effects overwhelmingly represent protein complex members (86% of top gene pairs; 23% expected by chance; Figure S1C). For example, the tuberous sclerosis complex proteins TSC1 and TSC2 represent the strongest coessential relationship in the genome (r=0.92, p=1e-269). Indeed, despite remaining highly statistically significant, the average magnitude of the top-ranked correlation for genes encoding transcription factors is substantially less than that for genes in the CORUM human protein complex database (Figure S1D). Moreover, in this network, non-informative clusters emerge which correspond to genes in the same chromosomal region, likely reflecting the confounding fitness effect of double-stranded breaks in regions with copy number alteration (CNA) (Aguirre et al., 2016) (Figure S1B). CNA-based corrections, now standard in defining gene essentiality from CRISPR-Cas9 screens (Meyers et al., 2017), address the magnitude of signal attributable to a gene at an amplified locus. However, this uniform correction does not address the shared patterns of relative essentiality between genes at that locus, fundamentally biasing correlation-based analyses. Thus, in a genome-scale, top-down coessentiality analysis, true biological signal is lost for genes encoding proteins which function independently from large molecular assemblies and/or have variable copy number across tumors – such as master regulator transcription factors of the cellular stress response (Figure S1E).

To address the limitations of existing coessentiality approaches, we developed a bottom-up approach to generate a locus-adjusted, rank-based network from central source node(s) of interest (Figure 1A). We first investigated HSF1, a transcription factor considered the master regulator of cytosolic proteostasis. Across cancer cell lines of diverse cell lineage and tumoral subtype, HSF1 loss negatively impacts cellular fitness, with a spectrum of relative essentiality within each lineage (Figure 1B). Notably, HSF1 is located on chromosome 8q, a region commonly amplified in cancer. In contrast to HSF1 transcript levels (Figure 1C), HSF1 essentiality was not correlated with copy number (Figure 1D). This reflects the copy number adjustment performed to obtain the CERES score, a continuous measure of the fitness effect of individual gene loss (Meyers et al., 2017). However, the genes most coessential with HSF1 still showed substantial enrichment for genes in the same chromosomal neighborhood (p = 7e-5; Figure 1E). We found that correcting for the median fitness effect of neighbor genes prior to performing genome-wide correlations eliminated this locus bias (Figure 1E).

**Figure 1:**
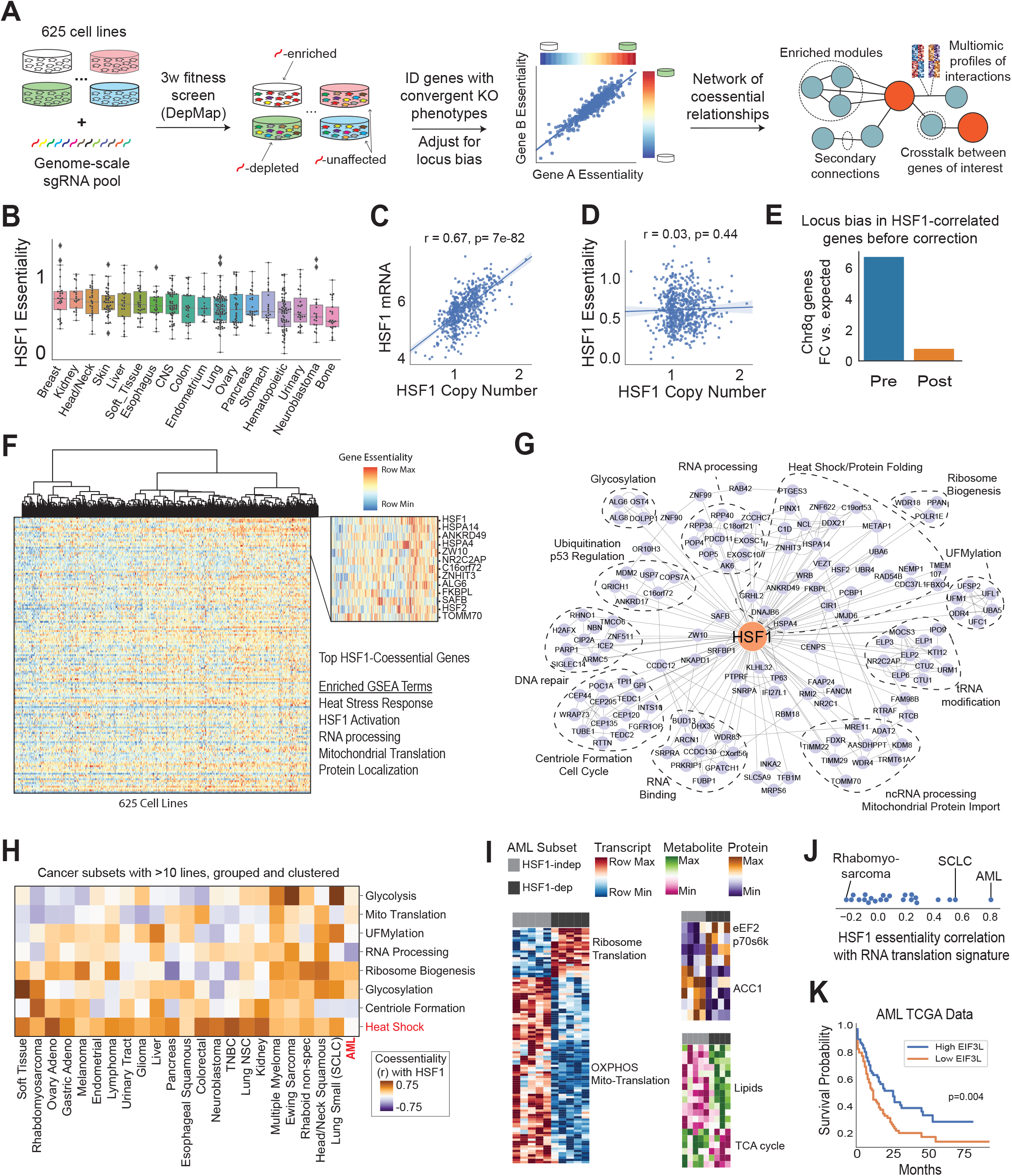
Developing a coessentiality network for the master regulator of cytosolic proteostasis. A. Schematic of overall approach. B. HSF1 loss is deleterious to cell fitness across lineages and has an essentiality spectrum within lineages. Essentiality is defined as the inverse of the CERES score, which quantitates the fitness cost associated with sgRNAs targeting a given gene in a given cell line; one is typical for a core fitness-essential gene and zero represents no fitness effect. C. HSF1 copy number is associated with HSF1 transcript level. D. The magnitude of HSF1 essentiality is not predicted by HSF1 copy number. E. The genes with the most similar fitness profile to HSF1 are highly enriched for genes in the same locus (chr8q). This bias is corrected by subtracting the median essentiality of 10 neighbor genes (5 upstream, 5 downstream) prior to performing genome-scale Pearson correlations. A sliding window of 10 genes most specifically reduced this locus bias. F. Visualization of the top 200 HSF1-coessential genes, where columns are cell lines, rows are genes, and hue corresponds to row-normalized gene essentiality. Also shown are gene sets for which the HSF1-coessential genes are enriched (FDR < 0.0001). G. Network visualization of the top 100 HSF1-correlated genes (primary nodes) plus the top 5 coessential genes for each primary node (secondary edges/nodes). Nodes are visualized if they have more than one connection in the network. H. Cancer subtype-specific coessentiality patterns for modules enriched in the pan-cancer HSF1 network, including a conserved heat shock correlation in all subtypes but acute myeloid leukemia (AML). I. Multiomic data integration to characterize the high-HSF1-dependence state in AML lines. J. HSF1 essentiality is highly correlated with the transcriptional signature of translation/protein synthesis genes in AML but not most other cancers. K. The translation-dominant RNA multiomic signature associated with HSF1-dependence in AML stratifies AML patient prognosis, as depicted by the top statistically-associated gene, EIF3L. P value from log-rank test.

We next profiled the top ranked HSF1-coessential genes using our locus-adjusted method. Remarkably, despite lower magnitudes of correlation as compared with protein complexes, the genes with fitness profiles most similar to HSF1 included genes having known functional relationships with HSF1, such as Heat Shock Factor 2 (HSF2), HSP70 and HSP110 family members (HSPA4, HSPA14), and HSP90 ligands (ANKRD49, FKBPL). Gene set enrichment analysis (GSEA) of top HSF1-coessential genes confirmed enrichment for canonical HSF1-related processes (i.e. chaperone-mediated protein folding; Figure 1F). GSEA also corroborated pathways with emerging relationships to HSF1, such as ribosome biogenesis and RNA processing, where HSF1 senses ribosome activity during translation (Santagata et al., 2013), and mitochondrial protein import, where HSF1 drives the multifaceted response to cytosolic accumulation of non-imported mitoproteins (Boos et al., 2019).

To better understand functional modules within the HSF1 coessential gene set, we generated an HSF1 coessentiality network containing the top 100 HSF1-coessential genes and up to 5 secondary connections per gene (Figure 1G). Like GSEA, the network approach reveals functional modules connecting to HSF1, with the additional advantage of illuminating the interplay within and between modules. For example, enzymes critical to protein glycosylation or UFMylation connect only to HSF1 and other genes in the same functional module. On the other hand, nucleolin (NCL) – a heat-shock responsive nucleolar protein which functions in the first step of rRNA processing (Daniely et al., 2002; Ginisty et al., 1998) – is central to a subnetwork which bridges the heat shock and RNA processing moedules.

We next investigated whether the HSF1 coessentiality network would vary by cancer subtypes, indicating lineage-specific biological relationships which did not emerge in pan-cancer analyses. As expected, the correlation of HSF1 with heat shock module genes was the most consistent correlation lineages, indicating that this canonical role of HSF1 is highly conserved across tissue types and genetic backgrounds (Figure 1H). Gene sets corresponding to other biological processes which have been previously linked to HSF1, such as glycolysis (Dai et al., 2007; Santagata et al., 2013; Zhao et al., 2009), had similar knockout phenotypes to HSF1 only in some cancer subsets, suggesting that context may drive these relationships (Figure 1H).

A notable exception to the otherwise subtype-conserved correlation between HSF1 and heat shock genes was found in acute myeloid leukemia (AML), where no relationship was observed between HSF1 and the heat shock module (Figure 1H). Rather, the most correlated module comprised genes involved in ribosome biogenesis. To better understand the contexts driving HSF1 essentiality in AML, we compared the transcriptome, metabolome, and proteome for the cell lines most dependent on HSF1 with the cell lines least dependent on HSF1. Strikingly, nearly every gene overexpressed in the HSF1-dependent AML lines encoded a protein involved in translation (enrichment p = 6e-25). Fittingly, protein levels of eukaryotic elongation factor 2 (EEF2) and p70s6k were elevated in these cell lines (Figure 1I), together indicating a high-translation phenotype which does not correspond to any known subtype of AML. Remarkably, a previous report described that inhibiting translation initiation with rocaglates potently inactivates HSF1, suppressing the high-translation malignant state in a manner which most potently impacted AML cells (Santagata et al., 2013). Notably, HSF1 dependence does not strongly correlate with this RNA signature of translation in other cancer subtypes (Figure 1J). Moreover, the observation that this phenotype exists in a spectrum within AML garners further consideration, given the ongoing exploration of rocaglates/translation initation inhibitors for cancer care (Cunningham et al., 2018). Gene expression data from AML patient tumors reveals that the protein synthesis genes identified in our cell line analysis above stratify patients into distinct prognostic populations (Figure 1K), suggesting that translation initiation inhibitors may be most efficacious in this patient subset. Moreover, these data suggest the existence of a low-translation, poorer prognosis group which may require alternative therapeutic approaches. Altogether, these data identify canonical and context-specific functions for HSF1, integrate genetic fitness data in a manner relevant to targeted cancer therapeutics, and serve as proof of principle for bottom-up, locus-adjusted coessentiality network analysis.

### A chaperone coessentiality network resolves functionally and spatially differentiated proteotoxic stress responses

While HSF1 is a critical upstream regulator of the proteotoxic stress response, the cellular proteostasis network also includes hundreds of effector chaperones and co-chaperones, many with distinct clients, patterns of activation, and subcellular localization. Reflecting the functional nature of chaperones to physically interact with client proteins, our understanding of chaperone biology is highly influenced by physical interaction studies, which are often constrained by the overexpression of tagged proteins, limited numbers of cell lines, and chemical inhibitors which fail to distinguish highly similar family members. Our genetic approach is not subject to these limitations and can identify functional relationships which do not require physical interaction. Thus, we expanded our network from HSF1 to the major chaperone families to provide functional insights into the broader cellular protein homeostasis network.

We first investigated the essentiality of the HSP40/J-domain protein (JDP), HSP70, HSP90, HSP110, HSP60/10, and small HSP chaperone family members (Figure 2A). Even within these families comprising highly related members, we observe great variability in the essentiality of individual chaperones. For example, within the HSP70 family, HSPA9 (Mortalin) and HSPA5 (BiP/Grp78) are broadly required for cellular proliferation (mean essentiality, 1.53 and 1.32; essentiality rank, 203 and 436 of 18333), whereas targeting either HSPA12A or HSPA12B has no deleterious effect on cellular fitness whatsoever (mean essentiality rank, 16738 and 13407). While most HSP70 family members are cytosolic, the two highly essential HSP70s represent the primary HSP70 in the mitochondria (HSPA9) and endoplasmic reticulum (HSPA5). Likewise, the ER-localized HSP90B1 (Grp94) is more essential than the cytosolic HSP90AA1 and HSP90AB1, suggesting that the specialization of these factors limits the capacity of the cell to buffer their loss via the expression of more closely related paralogs. Supporting this model, we observe that dependence on individual cytosolic HSP90 isoforms is predicted by relative expression of that isoform (Figure S2A,B). For example, leukemias preferentially express and depend on HSP90AB1, whereas cervical cancers preferentially express and depend on HSP90AA1 (Figure S2C). Because therapeutic modulation of chaperones remains a goal in many areas of patient care, these data argue for the consideration of paralog redundancy/buffering for any therapy targeted to a particular chaperone.

**Figure 2:**
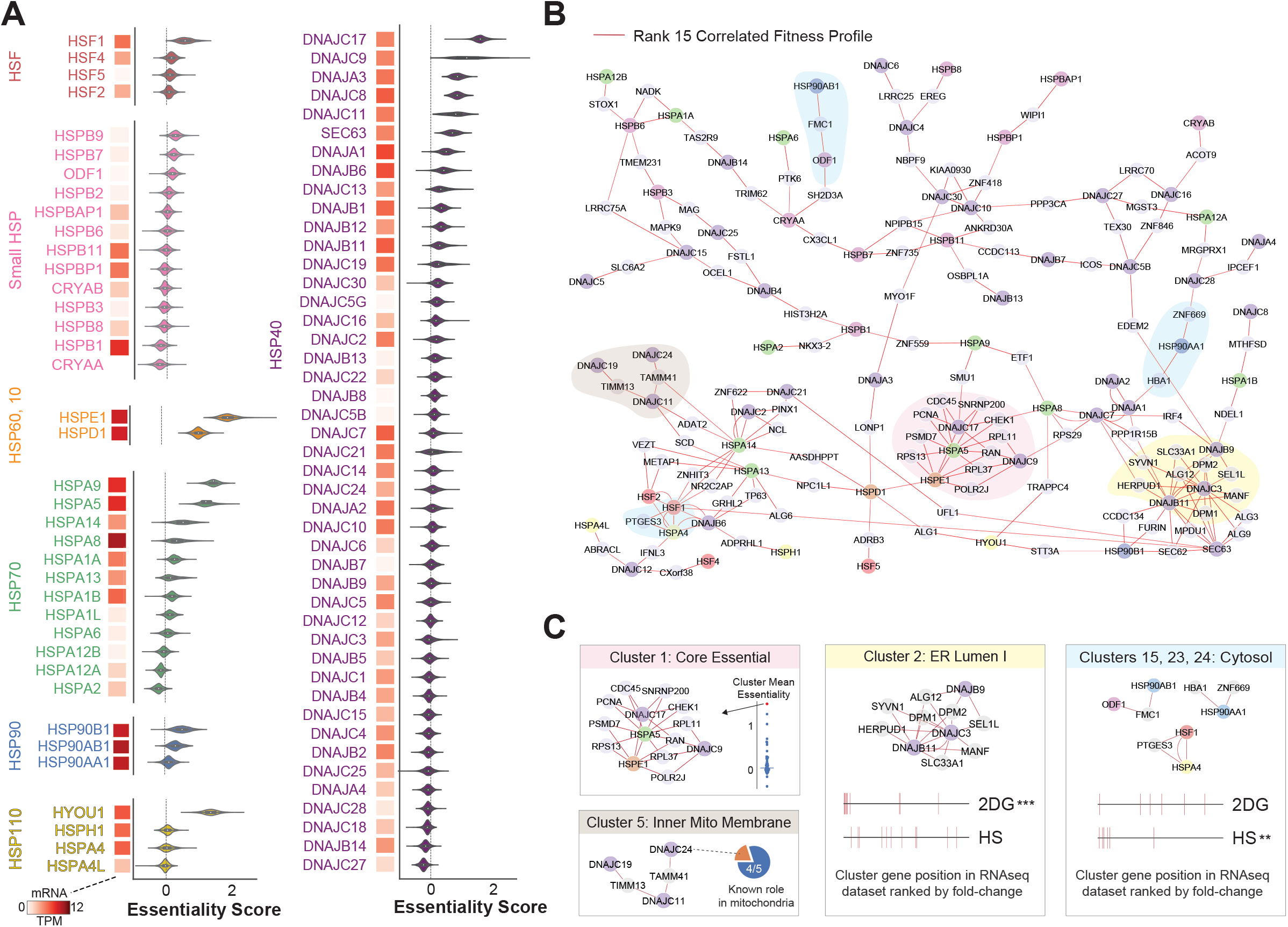
A chaperone coessentiality analysis reveals functionally and spatially differentiated networks. A. Essentiality scores and transcript expression of heat shock factors (HSFs) and major chaperone family members across 625 cancer cell lines. B. Rank 15 positive correlation coessentiality network of the genes in (A). Nodes are visualized if they have connections to more than one gene in the network. C. Selected Markov clusters from the network which represent biological functional groups corresponding to gene essentiality, localization, and/or divergent proteotoxic stress responses. RNAseq datasets derive from MDA-MB-231 cells treated with either 2-deoxyglusose (2DG) to induce ER stress or heat shock (HS; 42C × 1hr) to induce cytosolic proteotoxicity. Enrichment **p < 0.005, ***p<0.0005, Kolmogorov-Smirnov test.

Next, to investigate interplay between chaperone genes, we constructed a coessentiality network using these chaperone family members as source nodes (Figure 2B). Topologically, the chaperone network is more interconnected than would be expected by chance (p<0.001 vs. randomly permuted network), with several canonical relationships recapitulated. These included strong reciprocal connections between specific HSP70s and the JDP cochaperones with which they function, such as HSPA14 and DNAJC2. To further analyze the network, we performed Markov clustering, which revealed several distinct modules containing clear biological connections (Figure 2C). Cluster 1 contains the chaperones most correlated with core cellular fitness processes, such as DNA replication (PCNA), translation (RPL11, RPL37, RPS13), and transcription (POLR2J). Cluster 5 contained five member genes, four of which have known functions at the inner mitochondrial membrane, strongly suggesting that the outlier node (DNAJC24) also plays a role in mitochondrial protein maintenance. Remarkably, the 11 genes in Cluster 2 all localize to the ER lumen and have known functional connections to the ER stress response. On the other hand, multiple clusters emerged containing primarily to cytosolic proteins. To support the biological segregation of these cytosolic and ER lumen gene sets beyond localization, we investigated the transcriptional response of MDA-MB-231 cells to either 2-deoxyglucose (2DG; an inducer of protein misfolding in the ER lumen) or heat shock (HS; an inducer of protein misfolding in the cytosol). We found striking induction of ER cluster genes by 2DG, but not HS, and induction of cytosolic cluster genes by HS but not 2DG (Figure 2C, rightmost panels), indicating a functional differentiation of these modules in the response to distinct proteotoxic stressors. More broadly, these data demonstrate that bottom-up coessentiality network approach identifies functional modules, points of functional redundancy, and putative novel interactions within a highly interconnected and compartmentalized proteostasis network.

### Identifying topology and crosstalk between functional modules in a global stress response network

As disease states often impose multiple simultaneous stressors, we next sought to expand beyond proteostasis to investigate the crosstalk between diverse stress response programs. Thus, we used the approach outlined above to generate a global stress response coessentiality network centered around master regulators of the response to genotoxic, proteotoxic, hypoxic, nutrient, and oxidative stresses (Figure 3A). For this network, we also included highly-ranked negative correlations to capture antagonistic relationships (Figure 3B).

**Figure 3:**
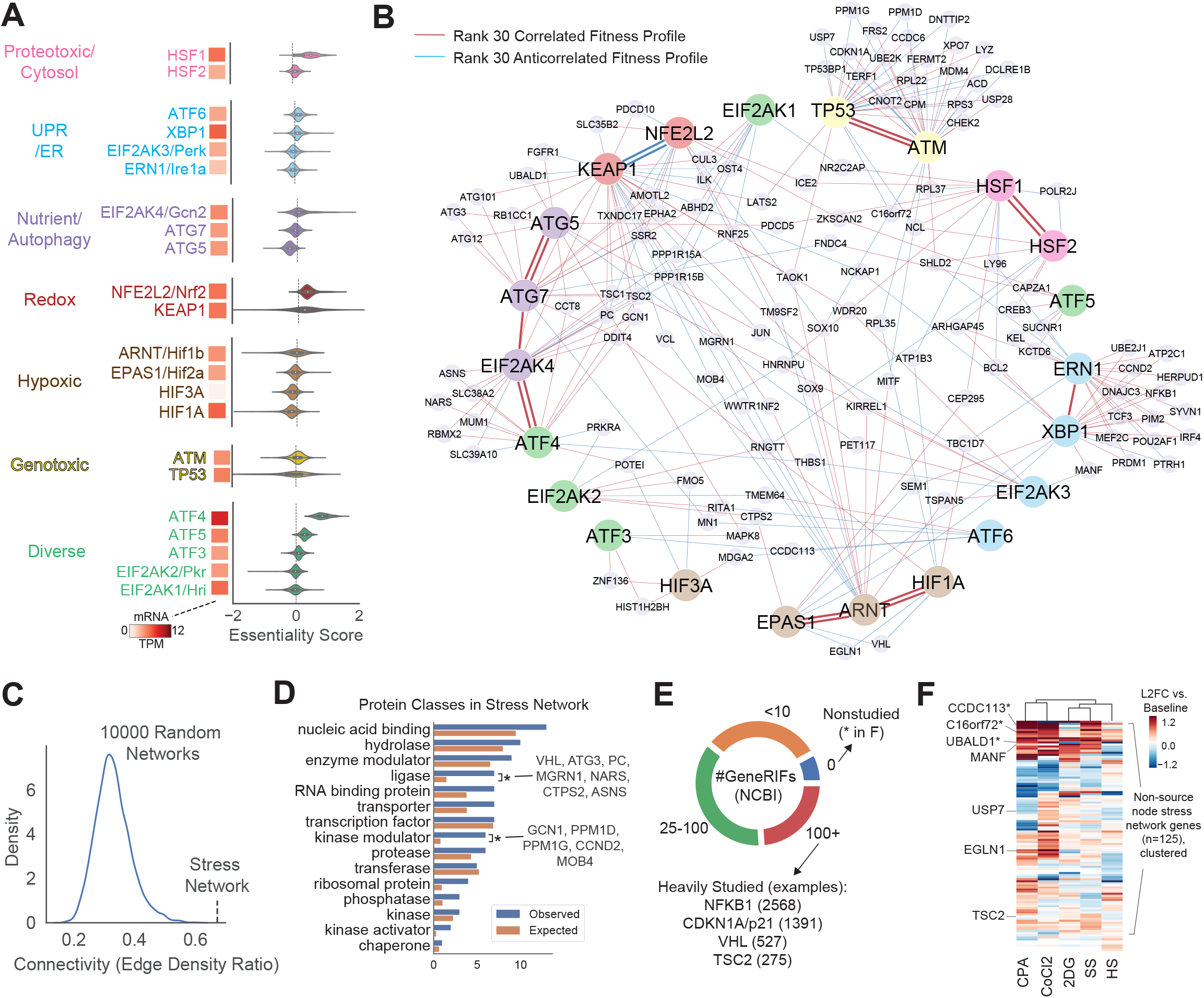
A global stress response coessentiality network. A. Essentiality scores and transcript expression of master regulators of diverse stress response programs across 625 cancer cell lines. B. Visualization of coessential gene relationships within stress response modules and between stress response modules at rank 30 for both positive and negative correlations. Stress networks thresholded at other ranks (10, 50, 100) are detailed in Figure S5. C. The stress network has greater connectivity (edge density ratio) than expected from 10,000 random network permutations using the same parameters. D. PANTHER functional classes of the 125 stress network genes pulled in as a function of coessentiality with source nodes; * indicates P<0.005, Fisher’s exact test. E. Representation of the degree to which stress network genes have been previously studied, using NCBI Gene References Into Function (GeneRIFs) as a surrogate for existing knowledge on a given gene. F. Transcriptome profiling by RNAseq of the stress network genes in MDA-MB-231 cells after treatment with Cyclophosphamide (CPA; genotoxic stressor), CoCl2 (hypoxia mimetic and oxidative stressor), 2-deoxyglucose (2DG; ER and nutrient stressor), heat shock (HS; 42C × 1hr; cytosolic proteotoxic stressor), or serum/amino acid starvation (SS; nutrient stressor). Selected genes elaborated in the text are labeled. *indicates gene with 0 GeneRIFs as indicated in (E)

The stress network (Figure 3B), comprising proteins with diverse molecular activities (Figure 3C), has far greater connectivity than would be expected by chance alone. Moreover, this network is replete with examples of canonical signaling relationships. For example, within the genotoxicity cluster, TP53 and ATM share connections with many genes critical to the response to double-stranded DNA breaks (e.g. CHEK2, CDKN1A/p21, TP53BP1). NFE2L2/NRF2 was highly anticorrelated with its canonical negative regulator, KEAP1, sharing multiple divergent nodes – i.e. nodes correlated to one source node and anticorrelated with the other. These divergent nodes included PDCD10/CCM3, which phosphorylates the Ezrin/Radixin/Moesin family proteins to promote fitness under oxidative stress (Fidalgo et al., 2012). The HIF family of transcription factors were also tightly connected, with the exception of the atypical HIF3A. Furthermore, VHL anticorrelated with both HIF1A and EPAS1 (HIF2A), but not ARNT (HIF1B), consistent with the known function of VHL to stimulate degradation of HIFα but not HIFβ proteins. The heat shock factor paralogs HSF1 and HSF2 were strongly connected, both displaying connections to the genotoxic stress cluster via genes such as NCL, which is known to mobilize from the nucleolus to nucleoplasm after heat shock or DNA damage in a p53-dependent process (Daniely et al., 2002).

Other clear modules in the stress network included a large cluster corresponding to ERN1 and XBP1. In response to proteotoxic stress in the ER lumen, ERN1 cleaves XBP1 to initiate one of the three branches of the unfolded protein response (UPR). Indeed, we find that ERN1 and XBP1 share a dense cluster of coessential genes enriched for UPR/ER stress signaling (e.g. HERPUD1, DNAJC3; enrichment p = 1e-9). Interestingly, EIF2AK3 and ATF6 – the sensors for the other two branches of the UPR – did not share many coessential genes with ERN1 and XBP1, consistent with recent data suggesting that specific contexts confer differential dependence on different branches of the UPR (Adamson et al., 2016). An exception to the lack of connectivity between UPR branches is MANF, which correlates with XBP1, but anticorrelates with EIF2AK3/PERK. Consistent with our network, MANF has conserved, divergent genetic interactions with XBP1 and EIF2AK3, but not ATF6 (Lindstrom et al., 2016). Moreover, MANF was recently identified to stabilize a subset of HSPA5-client complexes in the ER lumen (Yan et al., 2019). Indeed, HSPA5 is highly coessential with MANF in addition to correlating with XBP1 and anticorrelating with EIF2AK3, supporting a model in which MANF is a point of regulation between the functionally differentiated branches of the UPR.

Another highly interconnected cluster comprised genes involved in nutrient stress/autophagy, including the source nodes EIF2AK4 (GCN2) and GCN1 (nutrient sensors), ATG5 and ATG7 (critical genes for autophagosome formation), TSC1 and TSC2 (negative regulators of mTORC1), and ATF4. ATF4 is perhaps best known for its role downstream of EIF2AK3/PERK in the UPR, and does indeed connect to EIF2AK3 in the network via Thrombospondin 1 (THBS1), an ER-localized protein which promotes a protective ER stress response (Lynch et al., 2012). However, ATF4 clusters much more closely with the nutrient stress/autophagy genes, consistent with a more recently described role for ATF4 in mTORC1 signaling (Ben-Sahra et al., 2016). More globally, the autophagy cluster connected strongly with the oxidative stress module, featuring crosstalk genes such as TXNDC17 (TRP14), a disulfide reductase which has both peroxidase (Jeong et al., 2004) and autophagy-inducing (Tan et al., 2019; Zhang et al., 2015) activities.

While the stress network readily identifies canonical stress response signaling relationships, not all of the genes in the stress network have established connections to stress response biology. This may be because many of these genes have not been studied directly (Figure 3E). To test whether the genes in our network are regulated by stress, we treated MDA-MB-231 cells with five different perturbations which capture the breadth of the stress network (CoCl2: hypoxia mimetic, Cyclophosphamide/CPA: DNA damaging agent (CoCl2: hypoxia mimetic, Cyclophosphamide/CPA: DNA damaging agent, 2DG: ER stress inducer and metabolic stress, serum starvation (SS): nutrient stress, and HS: cytosolic proteotoxic stress). We observed that many genes in our stress network are stress-responsive at the transcript level, including three genes induced by multiple stressors which have not been studied directly (CCDC113, C16orf72, UBALD1; Figure 3F). While most genes were regulated by multiple stressors, certain genes corresponding precisely to one module were primarily induced by the stressor most associated with that module, such as TSC2 in serum/amino acid starvation, EGLN1 in CoCl2/pseudohypoxia, and the previously discussed MANF in 2DG/ER stress (Figure 3F). Altogether, the stress network recapitulates positive and negative regulatory interactions between distinct stress response modules, identifies genes which represent points of crosstalk between modules, and identifies a set of minimally-studied genes as putative members of an integrated stress response.

### Cancer cell dependence on dynamic stress modules reflects targetable vulnerabilities

While pan-cancer coessentiality analysis allows for the greatest sampling of distinct cell states, there are likely lineage-specific relationships within the stress network which might be obscured at the pan-cancer level. To identify such relationships, we constructed a stress response network using cancer cell lines from each lineage containing at least 10 cell lines and systematically assessed connectivity between source nodes (Figure 4A).

**Figure 4:**
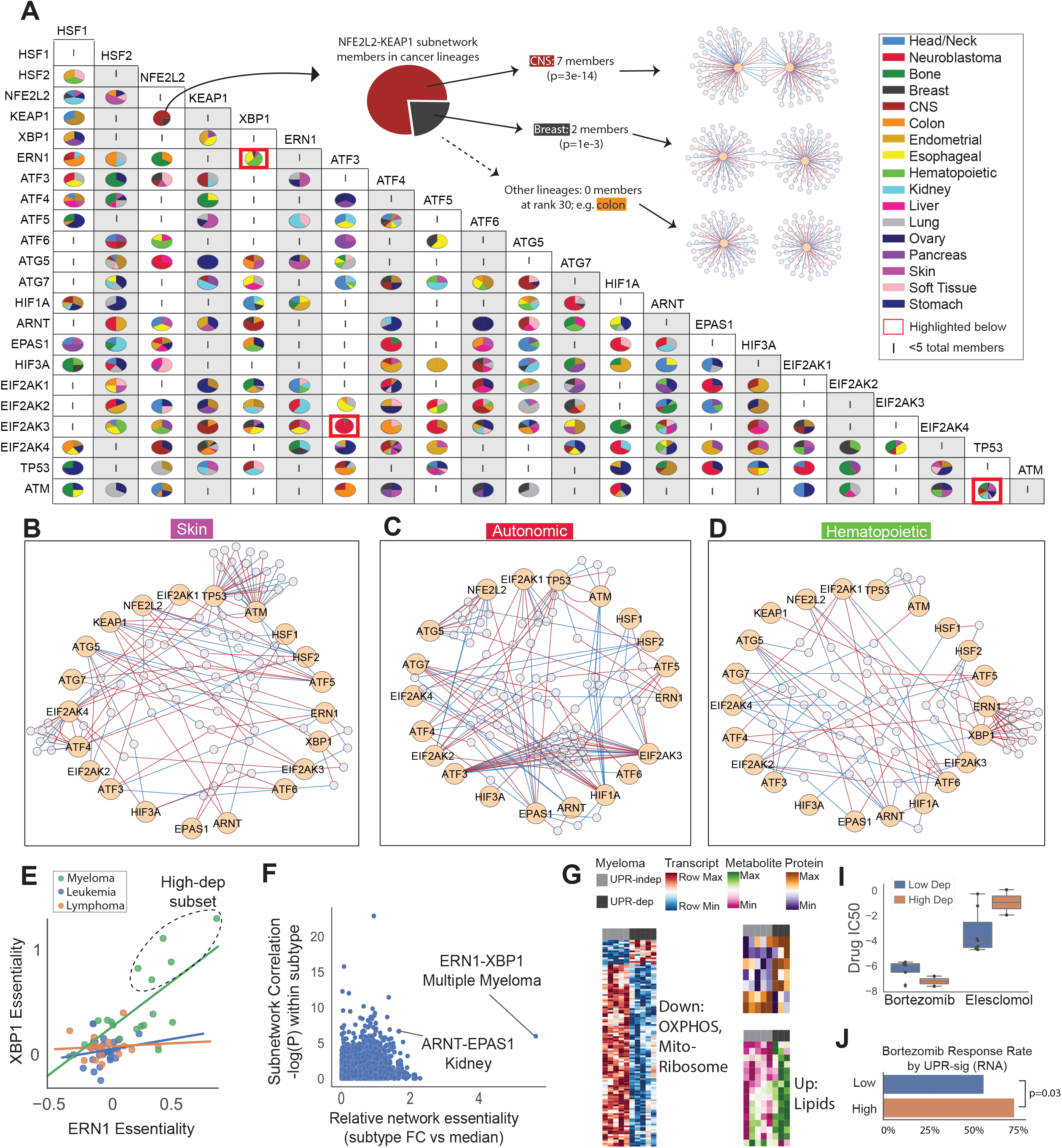
Systematic profiling of lineage-specific coessentiality patterns in the stress response. A. Lineage-specific subnetworks were created and profiled for relative enrichment of each possible source node pair subnetwork, with enrichment defined as membership relative to subnetwork membership across all other lineages and visualized as shares of a pie chart. B. The skin coessentiality network features the largest TP53-ATM genotoxicity network across cancer subsets, as well as crosstalk between all three branches of the UPR. C. The autonomic (neuroblastoma) network contains a highly anticorrelated network between ATF3 and EIF2AK3 which was not present in other lineages. D. The hematopoietic lineage features a striking ERN1-XBP1 subnetwork, but no relationships between other branches of the UPR (ATF6, EIF2AK3/PERK). E. Multiple myeloma drives the correlation of ERN1 and XBP1 in hematopoietic tumors, and there is a subset of myeloma particularly sensitive to the loss of ERN1 and XBP1. F. The tightly correlated ERN1-XBP1 network in multiple myeloma also represents a selective vulnerability. Other networks more moderately enriched for both essentiality and correlation included ARNT-EPAS1 (hypoxia) in kidney cancers. G. The myeloma lines most dependent on XBP1 and ERN1 have a low OXPHOS multiomic signature. I. The high-ERN1-XBP1-dependent myeloma lines are more susceptible to the proteasome inhibitor Bortezomib and substantially less sensitive to Elesclomol, which targets mitochondrial metabolism in myeloma lines resistant to proteasome inhibition (Tsvetkov et al., 2019). J. Patients enrolled in a clinical trial of Bortezomib (Mulligan et al., 2007) were more likely to respond if their tumors had >1 SD underexpression of the OXPHOS transcriptomic signature from (G). P values determined by Mann-Whitney U test.

Across lineages, the most highly connected stress response subnetwork corresponded to the TP53-ATM genotoxicity cluster, which had members in nearly every lineage, but was most predominant in skin cancers (18 members in skin; mean members in other lineages, 5.4) (Figure 4B). This is perhaps unsurprising, as melanoma classically contains the largest mutational burden of major cancers (Zhang et al., 2018). In contrast to the relatively conserved genotoxicity subnetwork, we identified an ATF3-EIF2AK3 network highly specific to neuroblastoma (17 members; one total member in other lineages). This network was comprised entirely of divergently correlated nodes – i.e. correlated with ATF3 and anticorrelated with EIF2AK3/PERK, or vice versa (Figure 4C). Notably, in neuroblastoma, ATF3 and EIF2AK3 are each the most anticorrelated gene for the other (r= −0.87, p=6e-6) while they are unrelated in the pan-cancer analysis (r=-0.04, p=0.3). ATF3 functions as a transcriptional repressor, and is induced by EIF2AK3 downstream of the ER stress response (Jiang et al., 2004). Querying ENCODE ChIP-seq data, we find that ATF3 can bind an EIF2AK3 enhancer (Figure S4). Taken together, these data suggest that ATF3 and EIF2AK3 operate in a negative feedback loop particularly relevant to neuroblastoma.

Another subnetwork with lineage-enriched membership was the ERN1-XBP1 UPR subnetwork, which had 15 members in the hematopoietic lineage network (mean membership in other lineages, 0.75) (Figure 4D). Interestingly, unlike the skin network (Figure 4B), which featured crosstalk between all three branches of the UPR, the hematopoietic lineage network was enriched only for the ERN1-XBP1 branch. We find that that this ERN1-XBP1 relationship in hematopoietic cells is primarily driven by multiple myeloma (Figure 4E), a malignancy characterized by constitutive UPR signaling due to high protein secretion load (Obeng et al., 2006). (Figure 4E). Strikingly, the XBP1-ERN1 network in myeloma was enriched for both network connectivity (highly correlated within the lineage) and network essentiality (selective essentiality within the lineage) (Figure 4F). Thus, the ERN1-XBP1 UPR branch is an exquisite vulnerability specific to a subset of multiple myeloma which could, in principle, be exploited by targeting any individual member of the network.

To further investigate this ERN1-XBP1-dependence in multiple myeloma, we compared available transcriptome, metabolomic, and drug sensitivity data for the myeloma lines most dependent on this network with those least dependent on the network. We found that this high-dependency subset of myeloma was characterized by a transcription and metabolic signature suggesting diminished oxidative metabolism (Figure 4G). Interestingly, these cell lines were more sensitive to Bortezomib (Figure 4I), a proteasome inhibitor used clinically in multiple myeloma which induces pro-apoptotic UPR signaling (Obeng et al., 2006). In contrast, the low-UPR-dependent, relatively proteasome inhibitor-resistant subset of myeloma was particularly sensitive to Elesclomol, an inhibitor of oxidative metabolism recently identified as a selective vulnerability in proteasome inhibitor-resistant myeloma (Tsvetkov et al., 2019). To investigate the relevance of these data to clinical care, we queried tumor microarray data from a clinical trial of Bortezomib, finding that the patients whose tumors shared this low-OXPHOS signature (>1 standard deviation from mean; n=29) were more likely to respond to Bortezomib therapy as compared with the other enrolled patients (n=159; Figure 4J). Altogether, these data demonstrate the utility of intra-lineage coessentiality network analysis to identify regulatory relationships and therapeutic vulnerabilities particularly relevant to cells of a specific lineage.

### C16orf72 is a stress-inducible modulator of the cellular adaptation to diverse stressors

To demonstrate the power of our approach to identify a novel gene critically relevant to the stress response, we chose to study C16orf72, a previously uncharacterized gene in our stress network which was also induced by multiple stressors (Figure 3C,E-F). Mining publicly available datasets, we found that C16orf72 is indeed expressed across cancer cell lines, normal tissue, and tumors (Figure S7), with its expression in tumors being prognostic in many cancers (Figure 5A). C16orf72 loss is broadly deleterious to the fitness of cancer cell lines particularly those derived from kidney tumors (Figure 5B). Evolutionarily, C16orf72 is highly conserved through C. elegans (Figure 5C), with CRISPR-Cas9 fitness screening data in Drosophila cells (Viswanatha et al., 2018) indicating that its importance for cellular viability is also conserved (Figure 5D).

**Figure 5:**
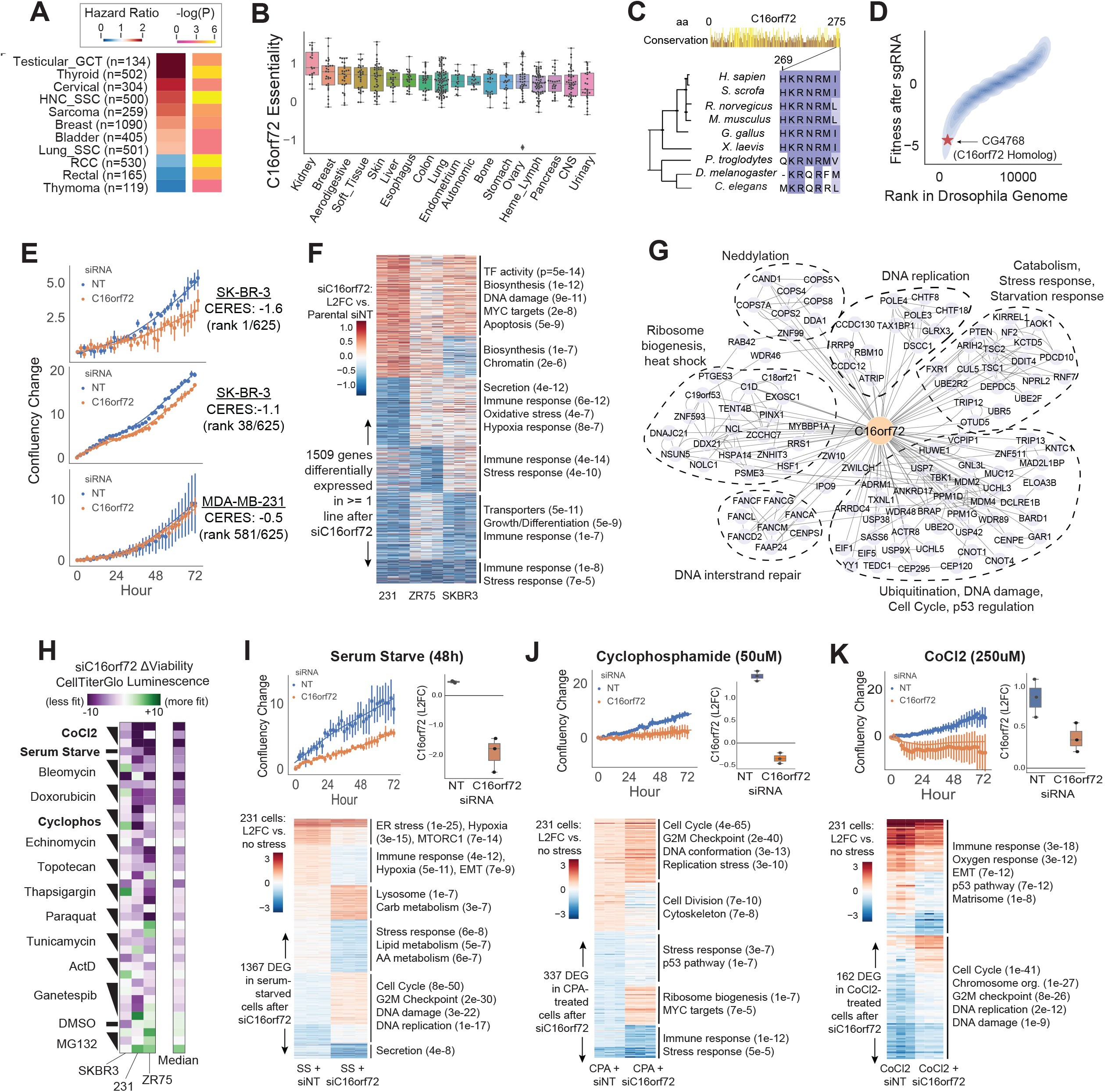
C16orf72 is a conserved and stress-induced modulator of diverse stress responses. A. C16orf72 expression is prognostic in many tumors. Subtypes shown had log-rank P value less than 0.05. B. C16orf72 loss is deleterious to the fitness of cancer cell lines across diverse backgrounds, particularly kidney cancer lines. C. C16orf72 is a 275aa protein conserved through C. elegans. Region shown represents a portion of a conserved nuclear localization signal as elaborated in Figure S6. D. A CRISPR screen in Drosophila cells (Viswanatha et al., 2018) indicates conservation of C16orf72’s importance to cellular proliferation. E. Validation of the Project Achilles fitness screen data suggesting that SKBR3 cell proliferation is highly dependent on C16orf72, ZR-75-1 cells are moderately dependent on C16orf72, and MDA-MB-231 cells are minimally affected by C16orf72 loss. F. Transcriptomic profiling of C16orf72 knockdown in these three cell lines identifies conserved regulation of pathways including multiple stress response networks. G. The C16orf72 coessentiality network demonstrates enrichment for ubiquitination pathways and multiple stress response networks. H. C16orf72 knockdown greatly hampers the ability of SKBR3, ZR75, and MDA-MB-231 cells to adapt to diverse stressors. I, J, K: Live cell imaging and RNA-seq of the three conditions with had the greatest combined effect on fitness in the background of C16orf72 depletion in MDA-MB-231 cells.

To validate the essentiality data for C16orf72 using an orthogonal approach, we tested the effect of C16orf72 depletion with siRNA on the short-term proliferation of three breast cancer cell lines with varying dependence on C16orf72: SK-BR-3 (C16orf72 essentiality score, 1.65; cell line rank, 1/625), ZR-75-1 (essentiality score, 1.09; cell line rank, 38/625), and MDA-MB-231 (essentiality score, 0.05; cell line rank 581/625). As expected, proliferation of SK-BR-3 cells was most impacted by C16orf72 depletion, followed by ZR-75-1, with no fitness effect at 72 hours in MDA-MB-231 cells (Figure 5E). Interestingly, C16orf72 depletion produced a similar overall transcriptional effect in each of these three cell lines (Figure 5F), consistently inducing genes related to DNA damage (e.g. CDKN1A/p21) and suppressing genes related to inflammatory, hypoxic, and other stress response programs.

C16orf72 emerged in our stress network, is induced by multiple stressors, and alters the transcription of stress response genes. Moreover, the C16orf72-coessentiality network contained clear modules related to genotoxic, hypoxic, proteotoxic, and nutrient stress (Figure 5G). To test the hypothesis that C16orf72 plays a broad role in the cellular response to stress, we designed a targeted screen to test the effects of C16orf72 depletion on cancer cells subjected to perturbagens which induce these diverse stress response pathways. We found that C16orf72-silenced cells were strikingly less tolerant to these stressors, except for the proteasome inhibitor MG132 (Figure 5H). Of note, MDA-MB-231 cells – which did not exhibit a proliferation defect upon C16orf72 depletion in the non-stressed, baseline condition – had a marked fitness defect when C16orf72 depletion was combined with stressors.

Because C16orf72 depletion most sensitized MDA-MB-231 cells to CoCl2 (hypoxia mimetic), serum starvation (nutrient stress), and CPA (DNA-damaging agent), we chose to further investigate the interplay between C16orf72 and these three stressors. Corroborating the ATP-based viability assay (Figure 5H), live cell imaging revealed reduced proliferation of C16orf72-depleted MDA-MB-231 cells in these conditions as compared with controls (Figure 5I-K). Moreover, knockdown of C16orf72, which was induced from baseline levels by each of the three stressors in the siNT control group, dramatically altered the transcriptional response to these stressors (Figure 5I-K). For example, 1367 genes were differentially expressed in serum starved cells without C16orf72 as compared to serum starved cells with C16orf72 (Figure 5I-K). Across all three stress conditions, knockdown of C16orf72 suppressed expression of genes corresponding to stress and inflammatory response pathways. Additionally, context-specific patterns of regulation emerged, whereby C16orf72 knockdown resulted in misregulation of amino acid metabolism genes under amino acid starvation, replication stress genes under interstrand crosslinking conditions, and oxygen response genes under hyperactive oxidative radical production. Altogether, our data identify C16orf72 as a conserved, stress-inducible gene which promotes cellular fitness in the face of diverse stressors.

### C16orf72/HAPSTR modulates the stress response by interacting with the stress-responsive E3 ligase HUWE1

To investigate the mechanism by which C16orf72 enables this striking resilience to stress, we first expressed a 3xFLAG-tagged C16orf72 construct in HEK293T cells. C16orf72 predominantly localized to the nucleus (Figure 6A), driven by a conserved nuclear localization signal at the protein’s N-terminus (Figure 5C, S6). Next, we performed affinity purification coupled with mass spectrometry to identify proteins which interact with C16orf72. The most prominent hit was the Hect ubiquitin E3 ligase HUWE1 (Figure 6B), which has known functional roles in the response to diverse stress conditions, such as genotoxicity, proteotoxicity, and hypoxia (Choe et al., 2016; Kao et al., 2018; Maghames et al., 2018). Confirming a physical interaction, we co-immunoprecipitated endogenous HUWE1 with C16orf72 using antibodies targeting both HUWE1 and FLAG in 293T cells (Figure 6C,D).

**Figure 6:**
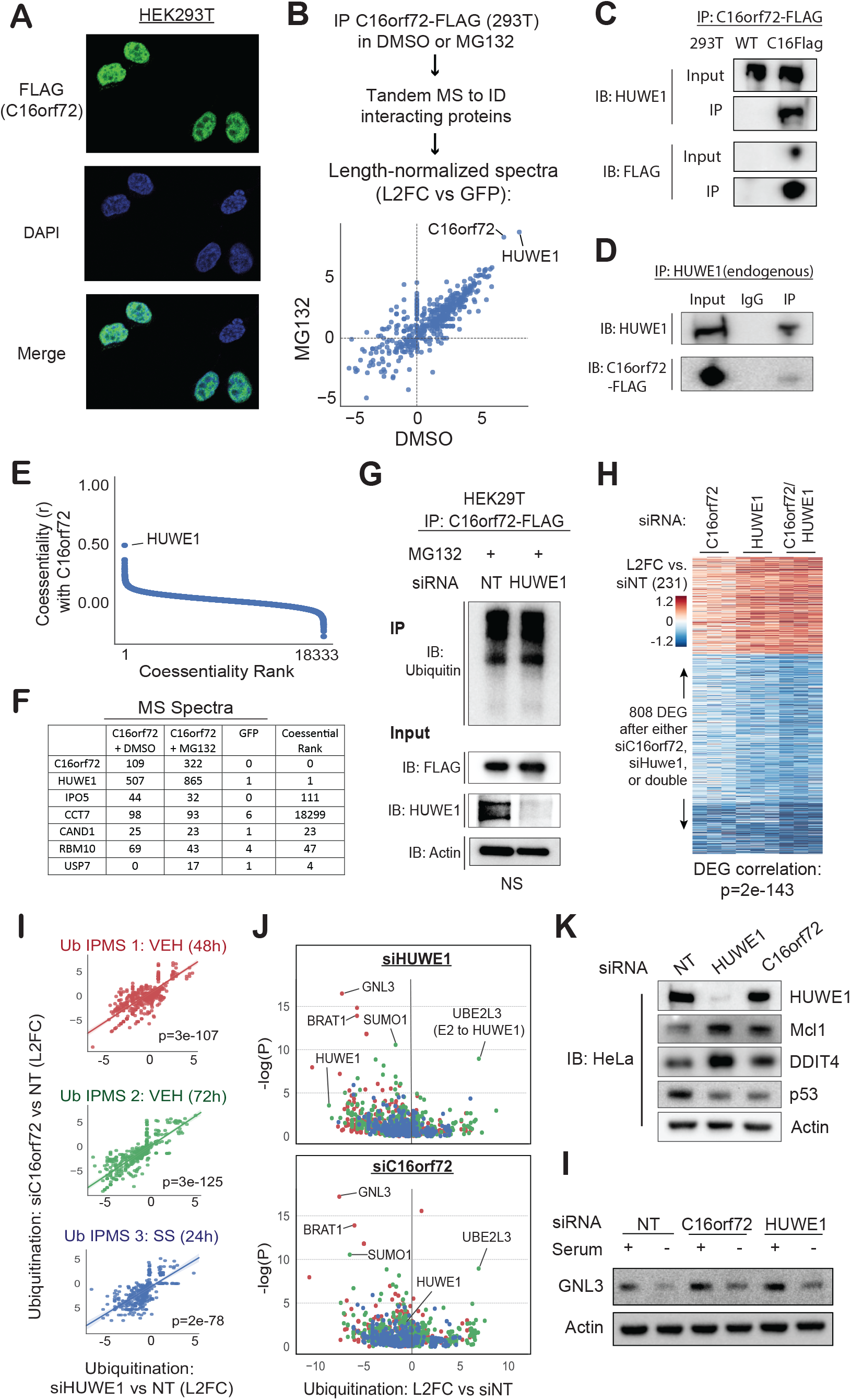
C16orf72 functions as a cooperative binding partner with the E3 ligase HUWE1. A. Expression of C16orf72-FLAG in HEK293T cells revealed a predominantly nuclear localization. B. C16orf72-FLAG affinity purification coupled with tandem mass spectrometry analysis identifies many binding partners for C16orf72, most prominently, the Hect E3 ligase HUWE1. Spectral counts were first normalized to the sum of spectral counts for the sample, then protein length, and finally compared vs. the maximum value for that protein from three separate control (GFP) IP-MS experiments. C. Immunoblot validation of HUWE1 IP using C16orf72-FLAG as bait. D. Co-IP of endogenous HUWE1 (bait) identifies C16orf72-FLAG. E. HUWE1 is the top-ranked gene in the C16orf72 coessentiality network. F. Many other proteins which interacted with C16orf72 were also coessential with C16orf72. G. C16orf72 protein levels and relative ubiquitination levels do not change with HUWE1 depletion, indicating that C16orf72 is not a ubiquitination substrate of HUWE1. H. Transcription profiling reveals that depletion of C16orf72, HUWE1, or both genes in MDA-MB-231 cells has a striking phenotypic overlap. I. The ubiquitinated proteome, as measured by FLAG-ubiquitin affinity purification coupled with mass spectrometry, is altered highly similarly by loss of either C16orf72 or HUWE1. J. C16orf72 and HUWE1 depletion converge on the under-ubiquitination of many proteins, such as GNL3/nucleostemin, and the over-ubiquitination of one protein, UBE2L3, which serves as an E2 for HUWE1. K. Representative immunoblots of established HUWE1 target protein levels after C16orf72 or HUWE1 siRNA treatment for 3 days. L. C16orf72 and HUWE1 depletion in HeLa cells increases baseline levels of GNL3/nucleostemin and reduce the capacity to eliminate the protein under stress (serum starvation).

Interestingly, HUWE1 is also C16orf72’s most coessential gene (r=0.49; Figure 6E). Several other proteins with which C16orf72 physically interacts also shared correlated fitness profiles with C16orf72 (Figure 6F). In contrast with the anticorrelated fitness profiles often observed between E3s and the targets they mark for degradation (e.g. MDM2 and TP53, r = −0.704), the highly convergent fitness profiles of C16orf72 and the E3 ligase HUWE1 suggest a cooperative function for these interacting proteins. Supporting the hypothesis that C16orf72 is not simply a HUWE1 substrate, we observe no significant change in C16orf72-FLAG expression or ubiquitination in response to HUWE1 knockdown (Figure 6G). Moreover, treatment of MDA-MB-231 cells with siRNAs targeting C16orf72, HUWE1, or both genes resulted in strikingly similar transcriptional profiles (Figure 6H). Of note, HUWE1 had a more substantial ma Taken together, these data strongly suggest a direct, cooperative physical interaction between C16orf72 and HUWE1.

To next address whether this putative cooperative function of C16orf72 may be related to the E3 ligase activity of HUWE1, we stably expressed FLAG-tagged ubiquitin under a doxycycline-inducible promoter in 293T cells and used FLAG affinity purification-mass spectrometry to identify changes in the ubiquitinated proteome after knockdown of either HUWE1 or C16orf72. Across three separate experiments, including baseline conditions (48h and 72h of knockdown) and 24h of serum starvation (72h of knockdown), C16orf72 and HUWE1 had nearly identical effects on the ubiquitinated proteome (Figure 6I). Specifically, C16orf72 and HUWE1 depletion independently resulted in under-ubiquitination of many proteins (Figure 6J). These included known HUWE1 substrates, such as TP53 (notably, C16orf72’s second-ranked fitness anticorrelation; r = −0.32, p = 6e-17), as well as putative HUWE1 substrates suggested from prior high throughput studies, such as GNL3/nucleostemin, a protein which promotes genome integrity during DNA replication (Tsai, 2014). Remarkably, of the 20 proteins which had highly significant changes in ubiquitination after C16orf72 and HUWE1 depletion, only UBE2L3 had increased ubiquitination. UBE2L3 is an E2 ligase which functions to transfer ubiquitin directly to the active-site residue of HUWE1 (a unique feature of Hect E3s). Together, increased ubiquitination of HUWE1’s E2 ligase and decreased ubiquitination of its substrates demonstrate that C16orf72 depletion is sufficient to markedly suppress HUWE1 ubiquitin ligase activity in a manner independent of a change in HUWE1 expression (Figure 6J,K). Moreover, C16orf72 depletion recapitulates the effect of HUWE1 loss on protein levels of established HUWE1 substrates (Mcl1, DDIT4, p53) as well as the putative substrate GNL3/nucleostemin (Figure 6K). Altogether, these data demonstrate a critical role for C16orf72 as a a stress-responsive cofactor for HUWE1, a fitness-essential gene known to modulate the stress response (Kao et al., 2018). Considering these data, we propose that this <u>H</u>uwe1-<u>a</u>ssociated modulator of <u>p</u>roliferation and the <u>st</u>ress <u>r</u>esponse be named HAPSTR. More broadly, the framework we use to identify HAPSTR provides a paradigm for the identification of other uncharacterized genes.

## DISCUSSION

Most genes in the human genome are not essential for cellular proliferation in standard culture conditions (Hart et al., 2015; Meyers et al., 2017; Tsherniak et al., 2017). Yet, many of these “nonessential” genes are critical to cell fitness in the context of physiological stress. Foundational work has demonstrated that, in yeast, only ∼17% of the genome is essential for growth in rich media (Winzeler et al., 1999), but nearly every gene (∼97%) becomes required for optimal growth in at least one chemical or environmental stress condition (Hillenmeyer et al., 2008). Here, we use the extraordinary heterogeneity of hundreds of cancer cell lines as a model for diverse cellular stress contexts, demonstrating that the contexts in which stress response genes become essential reflects the underlying biology of those factors. We exploit these context-specific essentiality patterns of known stress response genes to build a network of genes having similar context-specific fitness effects, effectively recapitulating canonical regulatory relationships, detailing treatment-relevant tumor dependencies, and identifying novel genes relevant to a global stress response network. Because an imbalance between stressors and the cellular capacity to adapt to those stressors underlies a broad spectrum of human disease, the data uncovered using this approach has broad therapeutic and prognostic implications.

The principle that convergent or epistatic knockout phenotypes may identify functional relationships between genes is not new (Dobzhansky, 1946). Great insights have been gained from pairwise genetic screens in model organisms, most prominently budding yeast (Costanzo et al., 2016), but the larger human genome has proved challenging for genome-wide combinatorial study of genetic perturbations. With the advent of high quality CRISPR-Cas9 screening libraries in recent years, the coessentiality approach has emerged as an alternative to pairwise genetic perturbations for the discovery of novel genes and genetic interactions (Boyle et al., 2018; Kim et al., 2019; Pan et al., 2018). These first coessentiality studies detail a top-down approach which effectively resolves major protein complexes and identifies functional clusters in the genome. Our locus-adjusted, rank-based approach addresses certain limitations of these studies, which include bias from copy number variable regions, overrepresentation of obligate protein complexes, and limited capacity to resolve genes which have dynamic functions across many cellular states – such as the transcription factors and chaperones of the cellular stress response. For example, HSF1 drives distinct transcriptional programs depending on cellular context (Filone et al., 2014; Mendillo et al., 2012; Scherz-Shouval et al., 2014), and both the chaperone (Joshi et al., 2018; Rizzolo et al., 2017; Rodina et al., 2016) and DNA damage response networks (Bandyopadhyay et al., 2010) are known to rewire under stress conditions.

In each of the networks elucidated in this study, canonical signaling relationships were recapitulated and modules of functionally related genes emerged. For example, within the chaperone network, we observe clusters distinguished both by subcellular localization and by patterns of induction in response to mechanistically distinct proteotoxic stresses. Of note, many chaperone and co-chaperone genes still have unresolved functional partners and localization, particularly within the HSP40/JDP family (Kampinga et al., 2019). Our network corroborates recent reports revealing the localization of individual JDPs, such as DNAJB11 in the ER lumen (Chen et al., 2017) and DNAJC11 at the inner mitochondrial membrane (Ioakeimidis et al., 2014). Thus, the novel relationships observed in our network, such as DNAJC24 at the inner mitochondrial membrane, are valuable targets for future study. Beyond chaperones, in the global stress network, we identify not only genes corresponding to individual stress pathways but also the genes which connect distinct stress programs. These genes, such as C16orf72/HAPSTR and NCL (bridging cytosolic proteostasis and DNA damage) and TSC1/2 and TXNDC17 (bridging oxidative stress and nutrient stress) represent points of crosstalk which likely play critical roles in diseased cells facing combinatorial stresses.

An overarching theme in these data is that cancer cells represent a breadth of disease-relevant cellular states which may offer functional insight not obtained from studies using limited numbers of cell lines or stress perturbations without a direct physiological correlate. For example, we identify a small subset (∼1%) of cancer cell lines particularly sensitive to the loss of genes involved in the ERN1-XBP1 branch of the UPR. Remarkably, these cell lines comprise multiple myeloma, a malignancy derived from plasma cells, which are characterized by a dramatically expanded ER network to accommodate the secretion of thousands of immunoglobulin molecules each second (Calame et al., 2003). Myeloma cells are burdened by a specific stress encountered in many disorders of secretory cells (e.g. pancreatic β cells in hereditary diabetes mellitus), and may thus serve as a more disease-relevant ER stress model than tool compounds such as thapsigargin (which induces ER stress by depleting ER calcium stores). Indeed, while all three branches of the UPR critically modulate the response to thapsigargin (Adamson et al., 2016), the ERN1-XBP1 branch is selectively required in these hypersecretory cells. In the clinical care of multiple myeloma patients, proteasome inhibitors are already employed to exploit the ER stress phenotype of malignant plasma cells. However, as the proteasome is a core essential component of all cells, our data suggest direct targeting of ERN1-XBP1 – for example, with available ERN1 inhibitors – may retain therapeutic efficacy while minimally affecting non-tumor cells.

Importantly, our data also indicate that there exists a spectrum of individual stress subnetwork utilization, even within a given lineage. Returning to the example of ERN1-XBP1, while this subnetwork was generally more essential in multiple myeloma, not all myeloma lines were equally impacted by ERN1-XBP1 loss. Thus, while targeting ERN1-XBP1 may have utility in the treatment of multiple myeloma, our data indicate that some tumors will be more susceptible than others to this modulation. Moreover, we find that the myeloma subset less likely to respond UPR modulation has a multiomic phenotype of increased oxidative metabolism and is, in turn, more susceptible to Elesclomol. Similarly, in AML, we observe variable dependence on the translation-HSF1 link, identify a transcriptional signature reflective of this phenotype, and demonstrate that this signature stratifies AML patients in a meaningful (Choe et al., 2016) fashion. It is worth noting that this approach to identify multiomic signatures of subnetwork dependency is unlikely to provide meaningful insight in pan-cancer analyses due to the powerful confounding effects of cell lineage. Thus, these data suggest a paradigm by which intra-lineage dependence on individual genes or gene networks can be translated into potentially actionable insight about an individual patient’s tumor.

Finally, we demonstrate the potential of our approach to identify novel genes relevant to a complicated biological network, such as C16orf72/HAPSTR. HAPSTR’s coessentiality network successfully predicted its cooperative relationship with HUWE1, as well as its role in the response to genotoxic, nutrient, proteotoxic, and oxidative stress. Corroborating our data, HAPSTR was also one of 117 genes identified to protect cells from inhibition of ATR, the primary sensor for single stranded DNA breaks, in a recently published screen (Hustedt et al., 2019). As HUWE1 is known to alleviate replication stress through an interaction with PCNA (Choe et al., 2016), our data reveal the likely mechanism of this observed chemical genetic interaction between HAPSTR and ATR inhibitors. Beyond genotoxicity, the interplay between HAPSTR, HUWE1, and proteins critical to diverse stress response programs sets the stage for many future studies. More broadly, because HAPSTR impacts the response to diverse stress states, is essential to some but not all cancer cells, and is prognostic of patient outcomes in diverse cancer diagnoses, HAPSTR may be a promising therapeutic target in cancer. It is also worth noting that HAPSTR is highly expressed in the developing brain (Figure S7B) and genomic alterations in both HAPSTR and its cooperative partner HUWE1 have been associated with neurodevelopmental disorders, such as autism (Bosshard et al., 2017; Levinson et al., 2011; Sanders et al., 2011). Thus, it is likely that misregulation of the evolutionally conserved, multi-stress-responsive protein HAPSTR plays a role in human pathology beyond neoplasia.

Far beyond HAPSTR and the response to stress, we emphasize that the approach detailed here is easily adapted to facilitate targeted study of other genes and pathways. To support such efforts, we make available an interactive web application (http://fireworks.mendillolab.org/) where individuals may input gene(s) of interest and quickly visualize a locus-adjusted coessentiality network. Moreover, this tool facilitates the integration of multiomic data to define the mechanisms underlying context-specific relationships. It is our hope that this resource will broadly augment efforts to identify functional interactions within biological networks.

## METHODS

### Curation of gene essentiality data in cancer cell lines

Data from CRISPR-Cas9 genome scale loss of function screening of 625 cancer cell lines was obtained from DepMap (https://depmap.org/portal/download/). CERES scores were used to quantify the fitness effect of individual gene loss, with “essentiality” in this paper represented as the inverse CERES score (i.e. more positive scores for genes which cause greater dropout of cells with guides targeting that gene. The screen performed in Drosophila melanogaster was identified through a search of BioORCs for any screens performed in Drosophila or C. elegans which targeted that species’ C16orf72 homolog.

### Integration and analysis of Cancer Cell Line Encyclopedia Multiomic Data

Processed RNA-seq, reverse phase protein array, copy number, and metabolomic data were obtained from the DepMap data portal (https://depmap.org/portal/download/). These data are described in detail in (Ghandi et al., 2019). For descriptive comparisons of different cell lines stratified by dependency signatures, the cell lines with 75^th^ percentile or higher dependency on that signature were compared with cell lines having 25^th^ percentile of lower dependency.

### Determination of coessential genes

To determine a locus-corrected coessentiality value for each gene pair, gene essentiality scores for each gene were subtracted from the median essentiality score for that gene’s nearest 10 neighbor genes (5 downstream, 5 upstream) on the same chromosome. Strength of coessential relationships are represented as Pearson r, with coessentiality rank used for all network analyses to mitigate bias from p-value inflation or differing numbers of cell lines in cancer cell line subsets.

### Coessentiality network visualization and clustering

Rank-based networks were constructed from a single or set of input genes, using a soft rank threshold for each analysis – i.e. correlations below the specific rank were not included. Edges are not weighted by correlation strength or rank. To remove potentially spurious correlations from genes only expressed in certain lineages, networks excluded transcripts where the median expression across cell lines was equal to 0. All networks were visualized in Cytoscape v3.7.2 (https://cytoscape.org/). Unless otherwise specified, networks used a force-directed layout with modest adjustments made by hand to improve legibility. The circular layout for the stress network was manually arranged following the groupings obtained by hierarchical clustering of source nodes. Statistical evaluation of network connectivity relative to randomly-permuted networks followed the method described in (Pan et al., 2018).

### Data analysis and visualization

Unless otherwise specified, data were analyzed with Python (version 3.6.4, Anaconda Inc.) using the modules Pandas (v0.23.4) and Numpy (v1.14.2) and visualization used the modules Matplotlib (v2.2.2) and Seaborn (v0.9.0). For heatmaps visualizing gene essentiality patterns across cell lines and cancer subsets, essentiality scores for a given gene are standard scaled to the minimum and maximum essentiality of that gene across all cancer cells. P-values were considered significant at an alpha of 0.05 or lower, as specified for each analysis. Benjamini-Hochberg false discovery rate correction was performed where indicated to adjust for multiple comparisons (Benjamini and Hochberg, 1995).

### Gene Set Enrichment Analysis

GSEA was performed using the Molecular Signature DataBase as accessible at http://software.broadinstitute.org/gsea/msigdb/annotate.jsp (Liberzon et al., 2015; Subramanian et al., 2005). The gene sets queried were as follows: hallmark (H), positional (C1), KEGG pathways (C2), REACTOME (C2), GO Biological Process (C5), and GO Molecular Function (C5).

### Protein conservation analysis

C16orf72 protein sequences for multiple species were obtained from ENSEMBL (http://useast.ensembl.org/index.html). Multi-sequence alignment was performed using Clustal Omega (https://www.ebi.ac.uk/Tools/msa/clustalo/), with visualization of the alignment in Jalview v2.11 (https://www.jalview.org/).

### Analysis of Patient Survival and Drug Response

For gene expression-stratified survival analysis, hazard ratios and log-rank test p-values were obtained through KM-plotter (https://kmplot.com/analysis/), using standard settings and the pan-cancer TCGA RNA-seq transcription dataset (Nagy et al., 2018). For AML analyses, we employed BloodSpot (http://servers.binf.ku.dk/bloodspot/) (Bagger et al., 2019). Bortezomib (Velcade) drug response likelihood was assessed using pre-treatment microarray data of multiple myeloma patients (GEO: GSE9782) (Mulligan et al., 2007).

### Functional classification and annotation of proteins

Grouping of proteins into functional classes was performed using PANTHER (pantherdb.org) to analyze a list of all genes targeted in the AVANA sgRNA library (Mi et al., 2019). NCBI Gene References Into Function (GeneRIFs; data downloaded from ftp://ftp.ncbi.nih.gov/gene/GeneRIF/), manually annotated blurbs summarizing findings in individual papers, were queried to bin the stress network genes by degree of prior existing knowledge.

### Experimental model and subject details

HEK293T and HeLa cells were grown in DMEM media supplemented with 10% FBS and 1% pen/strep. MDA-MB-231, ZR-75, and SK-BR3 cells were grown in RPMI media supplemented with 10% FBS and 1% pen/strep. Doxycycline inducible FLAG-Ub expressing HEK-293T cells were generated by transducing lentivirus containing pCW57.1-FLAG-Ub and selected in 2 μg/ml puromycin. All cells were passaged with Accumax unless otherwise specified.

### Plasmids, Lentivirus Generation and Infection

FLAG tagged ubiquitin was cloned in pCW57.1 vector (Addgene # 41393) between BsrG1 sites and verified through sequencing. PLenti6 vectors containing DNA constructs of interest were co-transfected with pMD2.G and psPAX2 into 293T cells using Lipofectamine 3000. After 48 hours, media was removed, filtered with an 0.45 µm filter, and centrifuged at 21000xg for 10 min to yield packaged lentivirus in the supernatant. Lentivirus was then added directly to cells for transduction. The C16orf72-FLAG overexpression vector contained a blasticidin resistance cassette, and stable overexpressing cell lines were selected for in blasticidin at 10 ug/mL (293T) or 20 ug/mL (MFC7, MDA-MB-231) for 5 days.

### Gene silencing and co-transfections

Smart-pool siRNAs were obtained for each target gene of interest, as well as a non-targeting sequences, and transfected using RNAimax (Thermo Fisher) using a standard protocol. Unless otherwise specified, cells were harvested 72h after siRNA transfection. Knockdown was confirmed for each siRNA experiment by qPCR or immunoblot.

### Drug and environmental perturbagen sensitivity in C16orf72-knockdown cells

MDA-MB-231, SKBR3, and ZR75 cells were reverse transfected with C16orf72 or non-targeting siRNA at 1000 cells/per well in a 384-well plate. After 1 day, RPMI media was replaced and drugs were arrayed at half-log intervals into wells using a Tecan D300E drug printer. RPMI media contained 10% FBS and 1% Pen/Strep, except for the serum starvation condition, which contained 0% FBS and 1% Pen/Strep. Drugs were selected based on pathways identified in the C16orf72 coessentiality network, as discussed. Drug concentrations, selected based on a pilot experiment to avoid complete cytotoxicity in the wild-type cell background, were as follows (µM): CoCl2 (25, 79, 250), Actinomycin D (ActD; 0.001, 0.0032, 0.01), Tunicamycin (0.05, 0.158, 0.5), echinomycin (0.01, 0.03, 0.1), Thapsigargin (0.05, 0.158, 0.5), Ganetespib (0.001, 0.0031, 0.01, 0.03, 0.1), MG132 (0.5, 0.158, 0.05), Topotecan (0.05, 0.158, 0.5), Doxorubicin (1e-6, 3.2e-6, 1e-5), Cyclophosphamide (0.05, 0.158, 0.5), Bleomycin (0.05, 0.158, 0.5), Paraquat (0.01, 0.032, 0.1).

### Proliferation and viability assays

Proliferation was assessed using live-cell images obtained via Incucyte. Cells were grown in 384-well plates, in triplicate for each condition, with whole-well images being taken every 2 hours. Area under the proliferation curve represented endpoint confluency subtracted from initial confluency for each well. Viability was assessed using an adapted CellTiterGlo (Promega) protocol; briefly, 384-well plates were brought to room temperature for 15 min, 12 uL of CellTiterGlo reagent mix was added to each well containing 50 uL of media, and the plate was agitated for 2 min. Luminescence was read out on a Tecan infinite M1000 pro platereader.

### Immunoblot

Protein samples were lysed in RIPA buffer or ubiquitin lysis buffer (as specified below) containing 1mM PMSF and a Roche Protease Inhibitor Cocktail tablet and passed through a 21 gauge syringe 15 times per sample. Protein concentration was assessed by standard BCA assay (Pierce, #23255), denatured in 4X Laemmli sample buffer containing beta-mercaptoethanol, and heated at 95C for 10 min. Electrophoresis was performed using 4-20% Bis-Tris gradient gels unless otherwise specified, with transfer to PVDF membranes using a 7-minute protocol on an iBlot machine. Membranes were blocked for 1 hour at room temperature in 5% fat-free milk. Primary and secondary antibodies were diluted in 5% fat free milk and exposed to membranes overnight at 4C and for 1 hour at room temperature, respectively. Imaging was performed with the BioRad ChemiDoc Touch Imaging System (732BR0783) after incubation for 2 min in HRP substrate (Immobilon, Millipore). Blots were analyzed using ImageLab v6.0.1 (BioRad).

### Immunofluorescence

Cells were grown on poly-D-lysine treated sterile coverslips in a 24 well-plate. Steps were performed at room temperature unless otherwise specified. Cells were washed three times with cold PBS, fixed with 4% paraformaldehyde for 10 min, and permeabilized with 0.2% Triton X100 for 5 min. Blocking encompassed incubation in 2% FBS for 30 min. Primary antibodies were diluted at 1:500 and secondary antibodies were diluted at 1:1000 in 2% FBS and exposed to cells for 1 hour each, with 3 PBST washes between. Coverslips were mounted to a slide using a DAPI/mounting mixture and allowed to dry overnight before imaging.

### Microscopy and Image Analysis

Images were acquired at 63x magnification using a Zeiss LSM800 confocal microscope. A z-stack slicing distance of 0.7uM was used, with final images visualized as an orthogonal projection of the maximal value per pixel across stacks. No non-linear adjustments were performed. ImageJ/Fiji (https://imagej.net/Fiji#Downloads) was used for all microscopy analyses.

### Immunoprecipitation

Cells were rinsed twice with cold PBS, removed from plates by scraping, and centrifuged for 4 min at 1000g and 4° C before pellet resuspension in cold lysis buffer. For immunoprecipitations of ubiquitin or of targets for downstream ubiquitination analysis, cells were vortexed and passed through a syringe in ubiquitin lysis buffer (2% SDS, 150mM NaCl, 10 mM Tris HCl pH 8, 1 Roche protease inhibitor tablet, 5mM N-ethylmaleide, and 1mM PMSF) and heated at 95° C for 10 min before dilution in ubiquitin lysis buffer with 1% triton and no SDS, resulting in a final SDS concentration of 0.2%. For all other immunoprecipitations, the lysis buffer was 1% NP40, 100 mM NaCl, 50 mM Tris pH 7.5, 0.2 mM EDTA, 5% glycerol, and 1mM PSMF and lysis was achieved by sonication in a 4C water bath (10 cycles of 30 sec on, 1 min off). After lysis, cells were spun at 21000xg and 4C for 10 minutes and the supernatant kept for input and immunoprecipitation. FLAG-immunoprecipitations were performed using M2 affinity agarose (Thermo Fisher).

### Mass Spectrometry

A protein gel band was submitted to the Northwestern University Proteomics Core Facility for an in-gel digestion. Peptides were analyzed by LC-MS/MS using a Dionex UltiMate 3000 Rapid Separation nanoLC coupled to a Orbitrap Elite Mass Spectrometer (Thermo Fisher Scientific Inc, San Jose, CA). Samples were loaded onto the trap column, which was 150 μm × 3 cm in-house packed with 3 um ReproSil-Pur® beads. The analytical column was a 75 um × 10.5 cm PicoChip column packed with 3 um ReproSil-Pur® beads (New Objective, Inc. Woburn, MA). The flow rate was kept at 300nL/min. Solvent A was 0.1% FA in water and Solvent B was 0.1% FA in ACN. The peptide was separated on a 120-min analytical gradient from 5% ACN/0.1% FA to 40% ACN/0.1% FA. MS^1^ scans were acquired from 400-2000m/z at 60,000 resolving power and automatic gain control (AGC) set to 1×10^6^. The 15 most abundant precursor ions in each MS^1^ scan were selected for fragmentation by collision-induced dissociation (CID) at 35% normalized collision energy in the ion trap. Previously selected ions were dynamically excluded from re-selection for 60 seconds.

Proteins were identified from the MS raw files using Mascot search engine (Matrix Science, London, UK; version 2.5.1). MS/MS spectra were searched against the UniProt Human database (SwissProt 2019, 20303 entries). All searches included carbamidomethyl cysteine as a fixed modification and oxidized Met, deamidated Asn and Gln, acetylated N-term as variable modifications. Three missed tryptic cleavages were allowed. The MS^1^ precursor mass tolerance was set to 10 ppm and the MS^2^ tolerance was set to 0.6 Da. The search result was visualized by Scaffold (version 4.9.0. Proteome Software, INC., Portland, OR). Peptide identifications were accepted if they could be established at greater than 90.0% probability by the Peptide Prophet algorithm (Keller et al., 2002) with Scaffold delta-mass correction. Protein identifications were accepted if they could be established at greater than 99.0% probability and contained at least 1 identified peptide. Protein probabilities were assigned by the Protein Prophet algorithm (Nesvizhskii et al., 2003). Proteins that contained similar peptides and could not be differentiated based on MS/MS analysis alone were grouped to satisfy the principles of parsimony.

### RNA-sequencing

RNA was extracted using a Qiagen RNeasy kit; briefly, cells were lysed in buffer RLT, nucleic acids were precipitated with ethanol and applied to columns, columns were treated with DNase, and RNA was eluted after washing/cleaning. Libraries were prepped using a QuantSeq 3′ mRNA-Seq Library Prep Kit FWD for Illumina (Lexogen) using 100ng of input RNA in an automated protocol adapted for the SciClone. Libraries were then analyzed for quality using the Agilent High Sensitivity DNA kit and for quantity using Qubit dsDNA HS assay, in 384-well format, using 20µL reactions in triplicate (19µL working reagent + 1µL sample or standard). For the Qubit assay, 11 standards were prepared from either 0-3ng/µL or 0-10ng/µL depending on BioAnalyzer concentrations. On plate reader, shake for 5 seconds, then read fluorescence: excitation: 480nm, emission: 530nm. Excitation/emission bandwidth: 5nm, settle time: 100ms. Sample concentrations were determined using the standard curve. Libraries were then pooled and sequenced using a NovaSeq 6000 SP Reagent Kit (100 cycles). Libraries were pooled at 25nM each, denatured with 1M NaOH added to a 0.2M final concentration (5 min at room temperature), and quenched with 200mM Tris HCl (pH 7). 1% PhyX spike-in (Illumina) was included. Pooled, denatured libraries were run on an Illumina NovaSeq using 51bp reads, 6bp index reads, and paired-end single read parameters.

## Data and Code availability

All code and data for these analyses will be deposited in public repositories. Our approach is implemented in in an interactive web application (http://fireworks.mendillolab.org/).

## Acknowledgements

We thank members of the Mendillo laboratory and Michael Drumm for comments on the manuscript. We thank Clara B. Peek for helpful discussions, Elizabeth Bartom for developing the Ceto pipeline used for RNA-seq analysis, and Kamil Slowikowski and Dan Wetherald for their advice on developing and deploying our web application. D.R.A. was supported by the NIH (5T32GM008152-33). M.L.M is supported by the National Cancer Institute of the NIH (R00CA175293) and the Susan G. Komen Foundation (CCR17488145). M.L.M was also supported by Kimmel Scholar (SKF-16-135) and Lynn Sage Scholar awards. Proteomics services were performed by the Northwestern Proteomics Core Facility, generously supported by NCI CCSG P30 CA060553 awarded to the Robert H Lurie Comprehensive Cancer Center, instrumentation award (S10OD025194) from NIH Office of Director, and the National Resource for Translational and Developmental Proteomics supported by P41 GM108569. D.R.F. was supported by the NIH (R01 GM111907).

## Author Contributions

Conceptualization: DRA and MLM. Data curation: DRA, JJ, BKC, YAG, NLK, MLM. Formal analysis: DRA, BKC. Funding acquisition: MLM. Investigation, DRA, KAM, DJA, RSS, SB, ST, BPO, BC, MLM. Methodology: DRA, JJ, KAM, SS, BKC, YAG, NLK, DRF, MLM. Project administration: MLM. Resources: MLM, NLK, SS, DRF. Software: DRA, JJ. Supervision: MLM. Validation: DRA, JJ, KAM, DJA. Visualization: DRA. Writing – original draft: DRA and MLM. Writing – review and editing: all authors.

---

**Figure S1:**
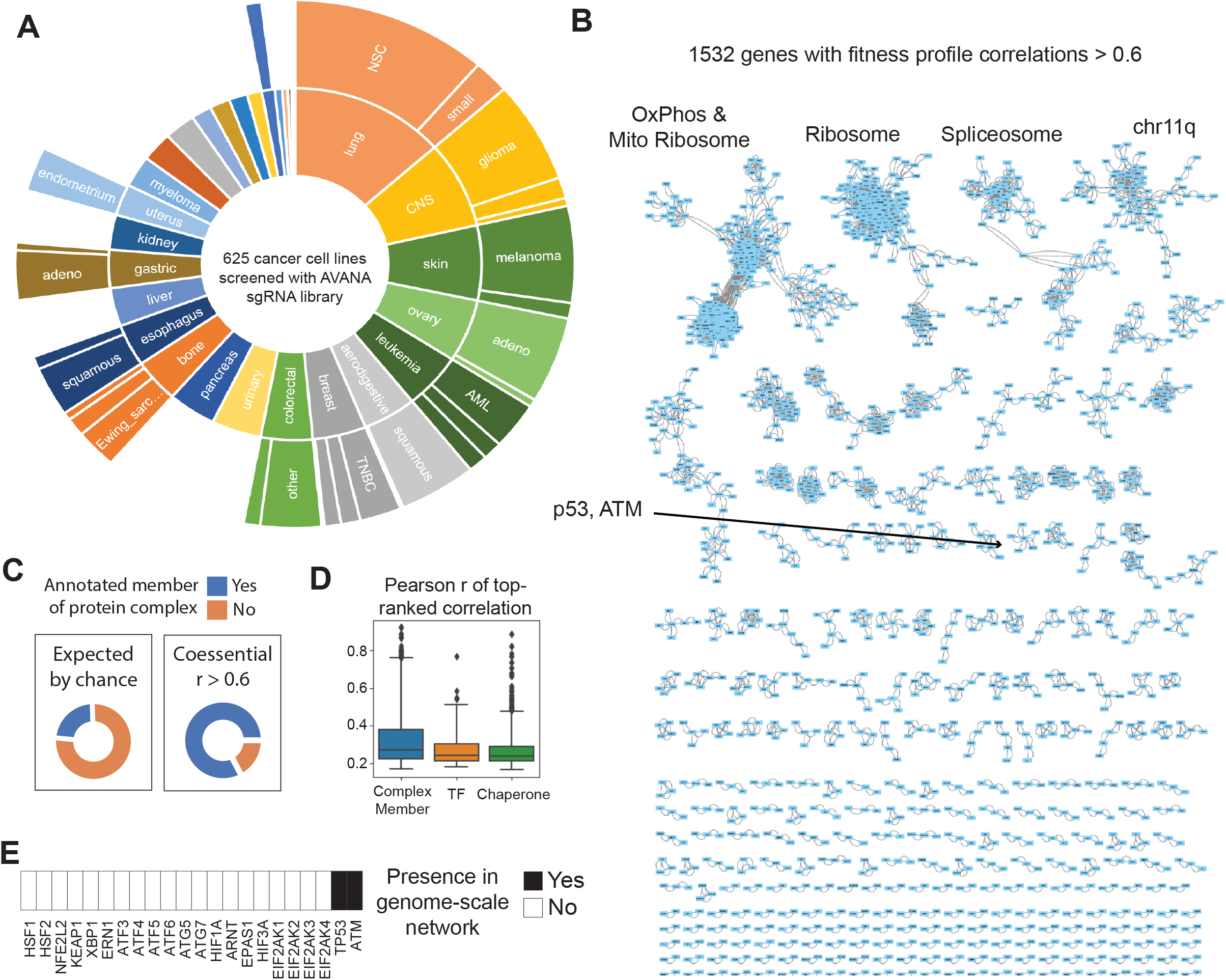
A top-down, genome-scale coessentiality network analysis does not resolve signal for major stress response genes. A. Schema representing the 625 cancer cell lines screened by the AVANA sgRNA library and included in these analyses. B. Visualization of the most coessential genes reveals dominant clusters corresponding to major molecular machines, as well as clusters corresponding to a known bias of assigning gene essentiality in CRISPR screens: copy number altered regions. C. Representation of proteins which form physical interaction complexes is substantially higher than would be expected in the genes which are most coessential across the genome D. The top fitness correlation for a generic transcription factor or chaperone is less than that of a protein complex member, suggesting reduced representation of these genes in any non-rank-based, top-down coessentiality approach. E. Of the 21 stress response master regulators of primary interest to this paper, only ATM and P53 have a functional module identified in a top-down genome scale clustering approach.

**Figure S2:**
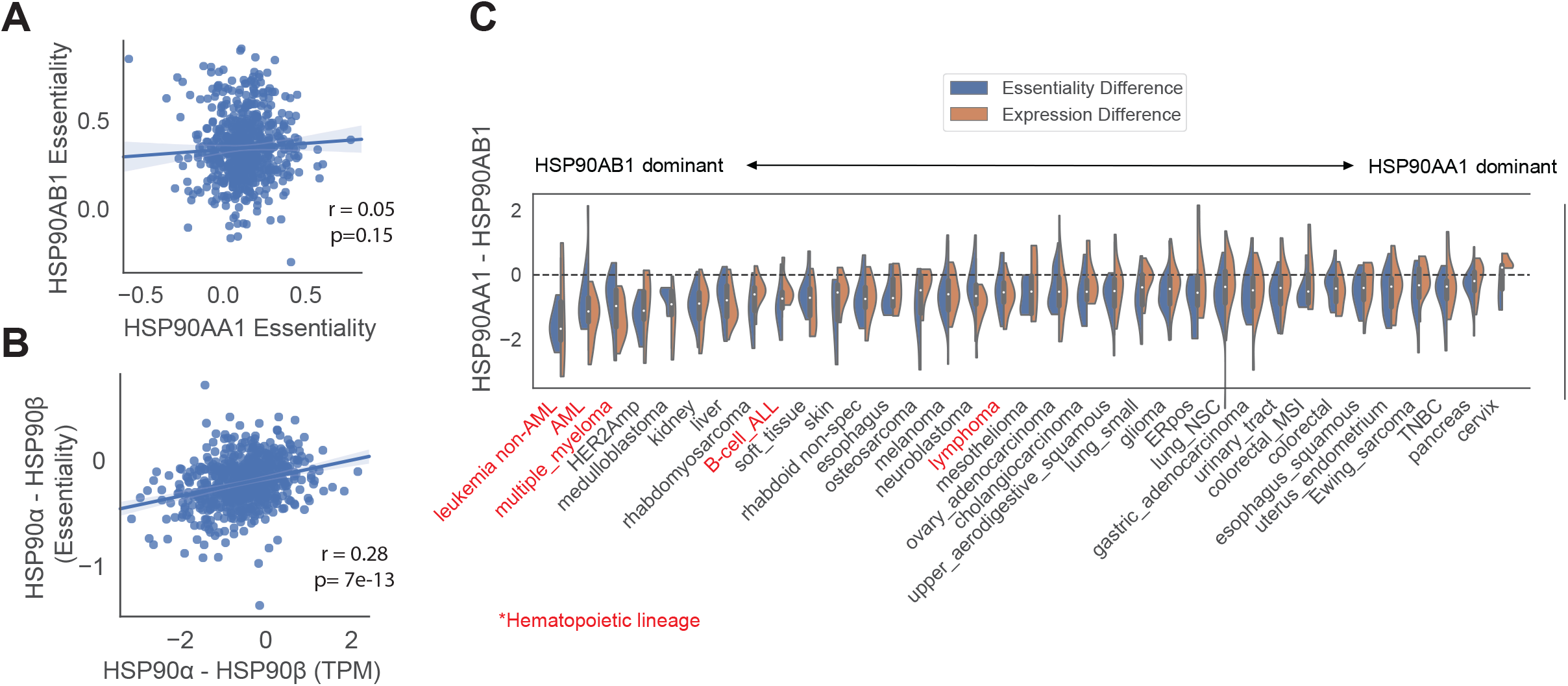
Cytosolic HSP90 isoform essentiality is predicted by HSP90 isoform expression in a manner related to cell lineage. A. HSP90AA1 and HSP90AB1 are not coessential, despite physically interacting in a major protein complex. B. Cytosolic HSP90 isoform expression predicts relative essentiality of that isoform. C. Certain tumor subtypes have higher expression and dependence on individual cytosolic HSP90 isoforms.

**Figure S3:**
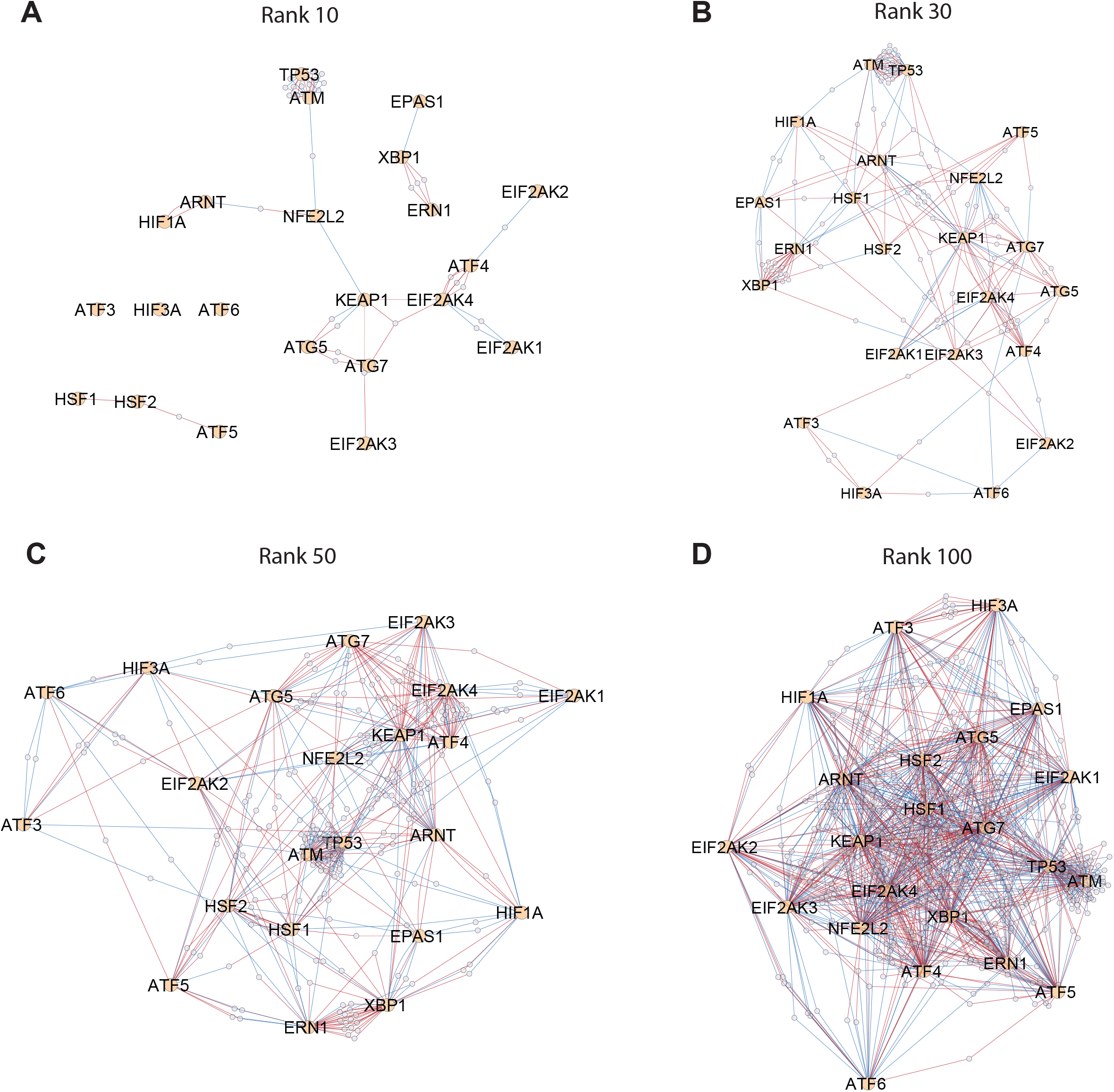
Alternative rank threshold coessentiality networks centered around stress response master regulators. Rank 10, 30 (as shown in Figure 3, but with hierarchical layout), 50, and 100 stress network organization using a force-directed layout. Red connections indicate positive correlations, blue indicate anticorrelations.

**Figure S4:**
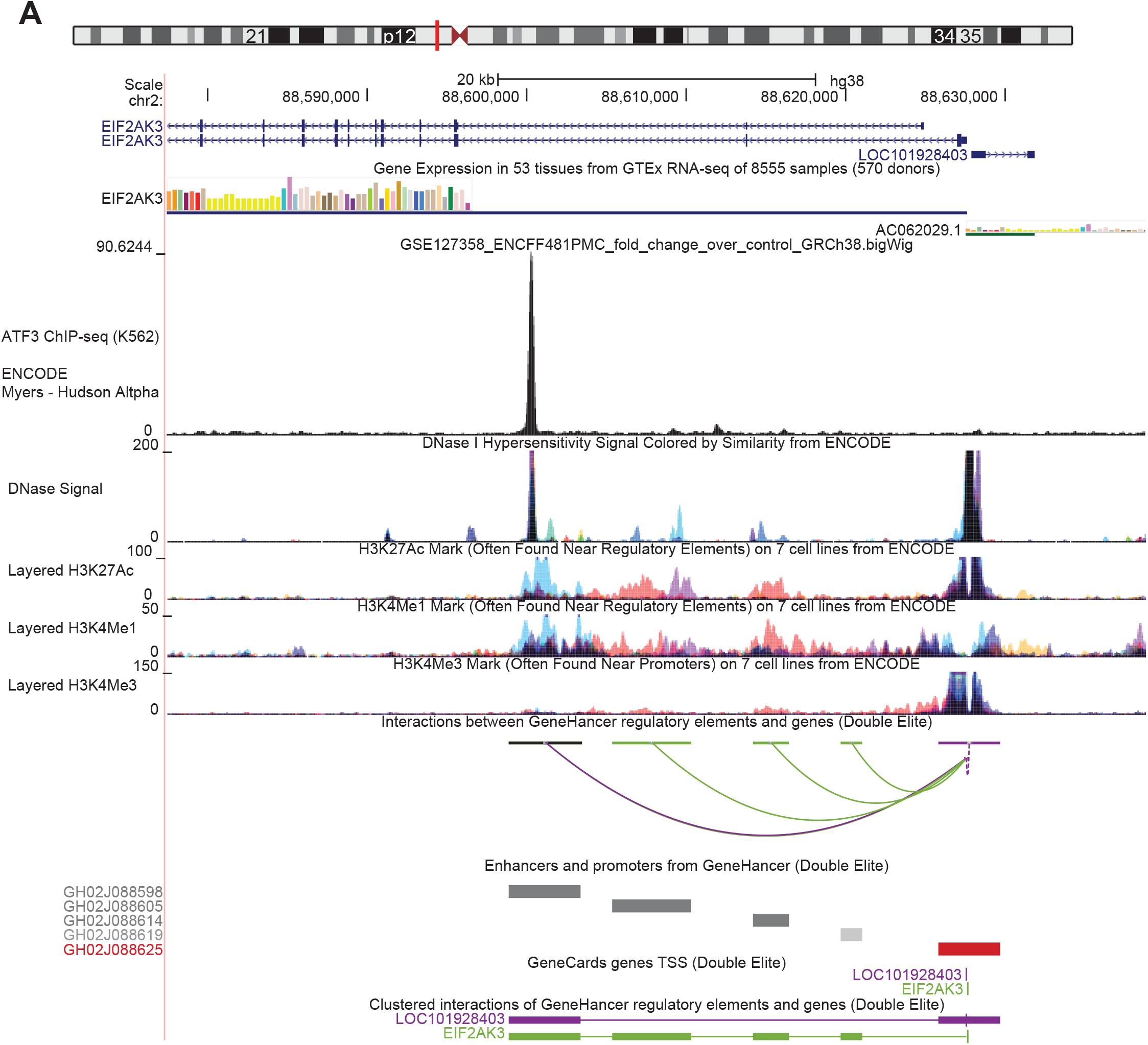
ATF3 binds to an EIF2AK3/PERK enhancer. ENCODE ChIP-seq data were queried for EIF2AK3, revealing that ATF3 binds strongly to an intronic regulatory element of EIF2AK3 cells characterized by high DNase sensitivity and annotated as an enhancer. Data shown derive from K562 cells.

**Figure S5:**
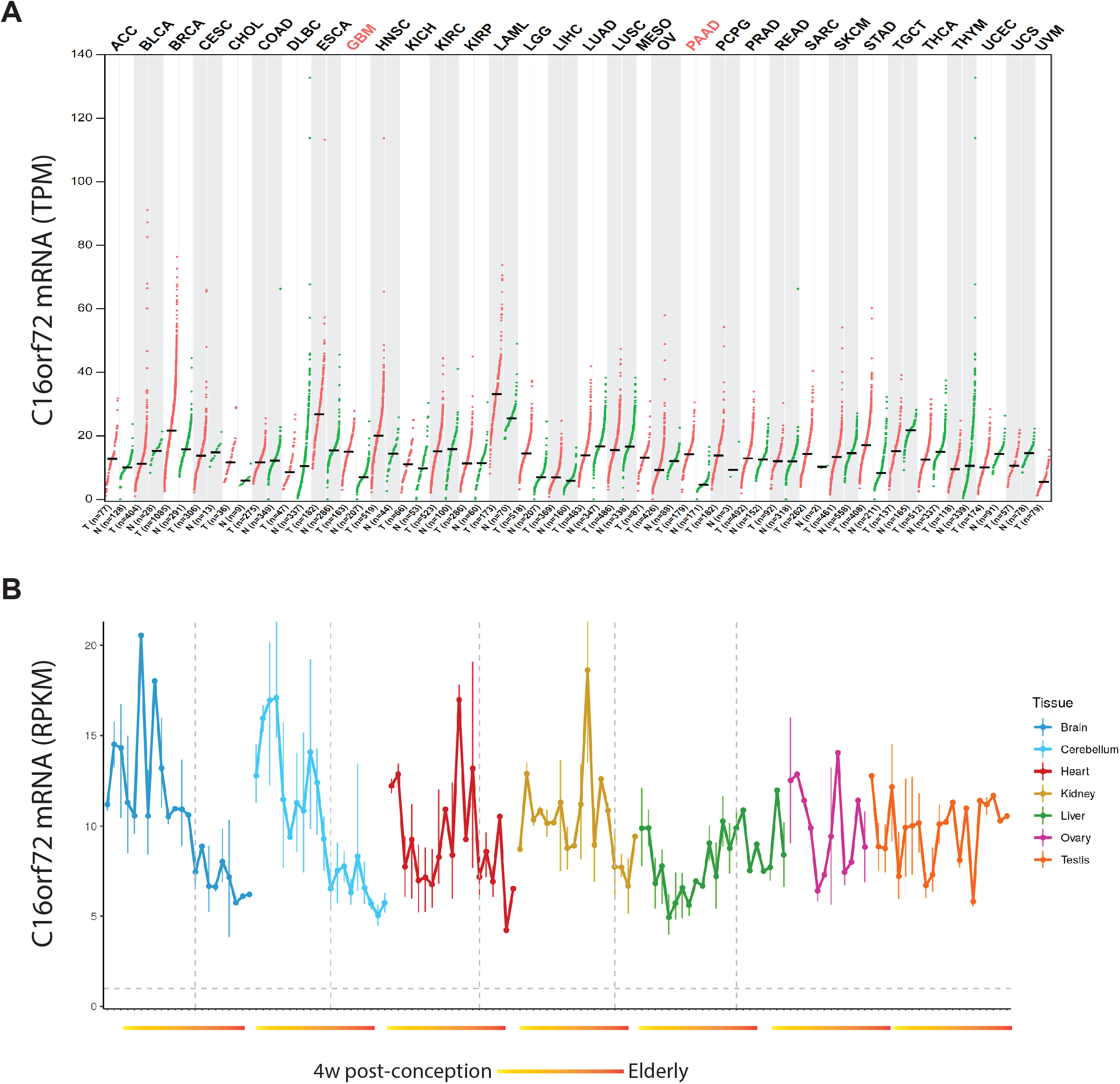
C16orf72 is expressed across broad cell types in adult tissues and tumors, as well as throughout human development. A. GEPIA output detailing tumor vs. normal transcriptional profiling of C16orf72; significant differences represented by red text. B. Expression of C16orf72 across human development; data from (Cardoso-Moreira et al., 2019).

**Figure S6:**
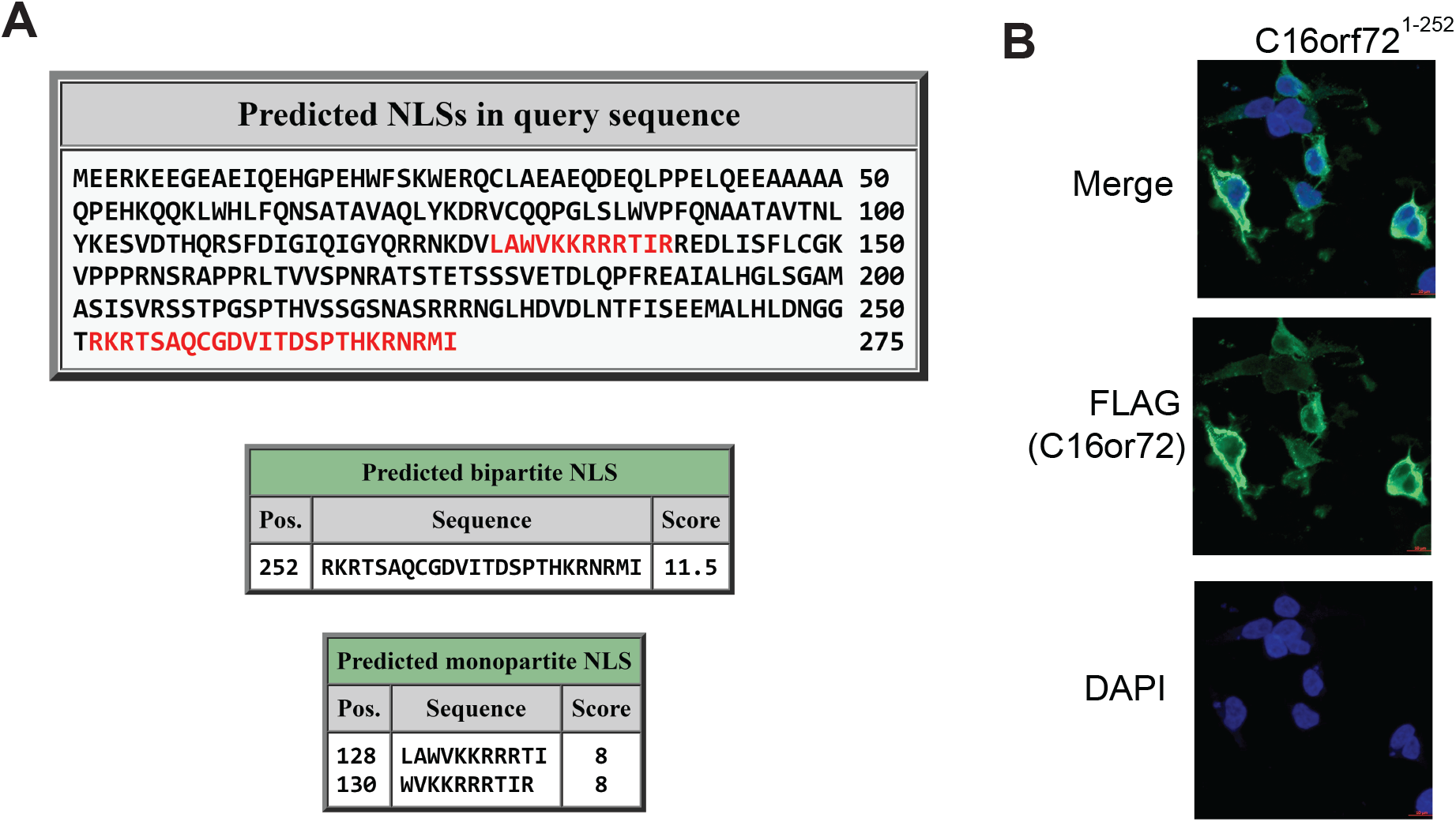
C16orf72’s nuclear localization is driven by an N-terminal bipartite NLS. A. Querying C16orf72’s amino acid sequence with cNLS mapper (http://nls-mapper.iab.keio.ac.jp/) revealed a predicted bipartite NLS at position 252-275. Score filter was set at 7, the most stringest cutoff. threshold, 7. B. Deletion of the putative NLS is sufficient to establish a cytosolic/perinuclear localization of C16orf72 in 293T cells as compared with its baseline nuclear localization (Figure 6A)

